# Checkpoint adaptation in repair-deficient cells drives aneuploidy and resistance to genotoxic agents

**DOI:** 10.1101/464685

**Authors:** Olga Vydzhak, Katharina Bender, Julia Klermund, Anke Busch, Stefanie Reimann, Brian Luke

## Abstract

Human cancers frequently harbour mutations in DNA repair genes, rendering the use of DNA damaging agents as an effective therapeutic intervention. As therapy-resistant cells often arise, it is important to better understand the molecular pathways that drive resistance in order to facilitate the eventual targeting of such processes. We employ repair-defective diploid yeast as a model to demonstrate that, in response to genotoxic challenges, nearly all cells eventually undergo checkpoint adaptation, resulting in the generation of aneuploid cells with whole chromosome losses that have acquired resistance to the initial genotoxic challenge. We demonstrate that adaptation inhibition, either pharmacologically, or genetically, drastically reduces the occurrence of resistant cells. Additionally, the aneuploid phenotypes of the resistant cells can be specifically targeted to induce cytotoxicity. We provide evidence that TORC1 inhibition with rapamycin, in combination with DNA damaging agents, can prevent both checkpoint adaptation and the continued growth of aneuploid resistant cells.

## Introduction

To preserve genome stability, eukaryotic cells have developed elaborate, and highly conserved, mechanisms to deal with DNA damage. In response to double-strand breaks (DSBs), a signalling cascade termed the DNA damage checkpoint (DDC) is activated to arrest the cell cycle before sister chromatid segregation occurs (reviewed in (Harrison and Haber, 2006)). The DDC is essential to prevent the propagation of cells with damaged chromosomes, which leads to genome instability, as well as providing time for the repair of the damaged DNA. The main steps in DDC activation in *Saccharomyces cerevisiae* comprise nucleolytic processing by the MRX complex and other nucleases (Lee et al., 1998; Mimitou and Symington, 2008; Zhu et al., 2008) to allow recognition of the damage site via Mec1/Ddc2 (Nakada et al., 2005). Subsequently, Mec1 activates the checkpoint effector kinases Rad53 and Chk1 utilizing the checkpoint mediator Rad9 (Ma et al., 2006; Sanchez et al., 1996; Weinert et al., 1994). To achieve a cell cycle arrest, Rad53 targets securin/Pds1 to prevent sister chromatid separation (Agarwal et al., 2003) and inhibits the pro-mitotic kinase Cdc5 (Cheng et al., 1998; Hu et al., 2001; Sanchez et al., 1999; Zhang et al., 2009), the yeast homologue of human Polo-like kinase 1 (PLK1) (Lee and Erikson, 1997). In addition, Rad53 triggers a transcriptional response to DNA damage (Edenberg et al., 2014; Gasch et al., 2001; Zhao et al., 2001).

Repair of DSBs is facilitated by two major pathways, namely homologous recombination (HR) and non-homologous end joining (NHEJ). The choice between these pathways is influenced by the cell cycle stage and hence the ability to resect the broken DNA ends, as well as the ploidy state (Mathiasen and Lisby, 2014). Diploid yeast cells actively suppress NHEJ throughout the cell cycle on the transcriptional level, rendering them dependent on HR to repair DSBs (Frank-Vaillant and Marcand, 2001; Kegel et al., 2001; Valencia et al., 2001). To engage in HR, DSB ends are resected and rapidly bound by the ssDNA binding protein complex RPA (Mimitou and Symington, 2008; Sugiyama et al., 1997). Rad52, which is essential for HR-mediated repair in yeast, then promotes the replacement of RPA with the recombinase Rad51 (Sugiyama and Kowalczykowski, 2002; Sung, 1997a). Rad52-mediated loading of Rad51 is assisted by Rad55 and Rad57 (Sung, 1997b). The Rad51-ssDNA nucleoprotein filament then scans the genome for intact homologous template sequences to drive the error-free repair reaction (Bernstein et al., 2011; Godin et al., 2013), (Mine-Hattab and Rothstein, 2013). Following successful DNA repair, the checkpoint is eventually extinguished and cells are able to continually divide, a process referred to as checkpoint recovery (Leroy et al., 2003; Pellicioli et al., 2001).

In certain scenarios, DNA damage can be considered irreparable as dictated by the nature of damage, i.e. where the damage occurs (e.g. telomeres), or the genetic context of the cell (e.g. loss of *RAD52* in diploid yeast). In response to persistent damage, DDC signalling can however still be terminated, in a process termed checkpoint adaptation (Sandell and Zakian, 1993; Toczyski et al., 1997). Checkpoint adaptation is considered to be a last effort to maintain cell viability and avoid cell death by attempting to either tolerate the DNA damage load, or repair by alternative means (Clemenson and Marsolier-Kergoat, 2009). In response to genotoxic stress and dysfunctional telomeres, checkpoint adaptation likely precedes genomic instability (Coutelier et al., 2018; Galgoczy and Toczyski, 2001) and facilitates the use of mutagenic repair pathways including break-induced replication (BIR) (Galgoczy and Toczyski, 2001). Importantly, checkpoint adaptation has been documented in both uni- and multicellular systems including budding yeast, *Xenopus* egg extracts, plant and cancer cells (Carballo et al., 2006; Kubara et al., 2012; Sandell and Zakian, 1993; Swift and Golsteyn, 2016; Syljuasen et al., 2006; Toczyski et al., 1997; Yoo et al., 2004). One striking similarity in the process of checkpoint adaptation across all organisms is the critical contribution of PLK1 (Liang et al., 2014) and its homologues such as budding yeast Cdc5 (Syljuasen et al., 2006; Toczyski et al., 1997). An allele of *CDC5* (*cdc5-ad*), which harbours a single point mutation (L251W), is specifically defective in checkpoint adaptation (Toczyski et al., 1997). The nutritional status of a cell can also impact the decision to undergo checkpoint adaptation (Klermund et al., 2014). More specifically, inhibition of the highly conserved TOR complex 1 (TORC1) by rapamycin treatment was shown to prevent checkpoint adaptation in response to chronic dysfunctional telomeres (Klermund et al., 2014). TORC1 not only promotes checkpoint adaptation in the presence of unrepaired DNA damage (Klermund et al., 2014) but also enables chromosome mis-segregation by activating the Glc7/PP1 phosphatase, an antagonist of the spindle assembly checkpoint (SAC) protein, Ipl1 (Tatchell et al., 2011).

It has long been recognized that yeast cells lacking Rad52 experience extensive chromosome loss after irradiation (Mortimer et al., 1981), however direct links to checkpoint adaptation have not been drawn. Extensive studies on yeast and mammalian cells with abnormal chromosome numbers (aneuploid cells) have identified a common aneuploidy-associated phenotype characterized by increased levels of proteotoxic stress resulting from imbalanced protein expression (Donnelly et al., 2014; Oromendia et al., 2012; Tang et al., 2011; Torres et al., 2010; Torres et al., 2007), delayed cell cycle progression (Beach et al., 2017; Stingele et al., 2012; Thorburn et al., 2013), metabolic stress (Tang et al., 2011; Torres et al., 2007), non-genetic heterogeneity between clones (Beach et al., 2017), hypo-osmotic-like stress responses (Tsai et al., 2019), as well as increased and persistent DNA damage (Blank et al., 2015; Passerini et al., 2016). Many of these phenotypes that exist in aneuploids are susceptible to being targeted by pharmacological agents (Tang et al., 2011).

In this study, we have used repair-defective diploid yeast as a model to better understand the consequences of checkpoint adaptation following a challenge with clinically relevant genotoxic agents. We observe that the re-growth of repair-defective cells almost exclusively depends on checkpoint adaptation. All of the adapted cells that we characterized had undergone chromosome loss, rendering them aneuploid, and had acquired resistance to further DNA damaging agents. By combining genotoxins with inhibitors of checkpoint adaptation or with aneuploidy-selective agents, we were able to synergistically enhance the cytotoxicity of repair-defective cells, while repair-proficient cells remained unscathed. These results suggest that combination therapies targeting checkpoint adaptation and aneuploidy-associated phenotypes together with conventional DNA damaging chemotherapeutics, may be considered in human cancers with mutations in DNA repair genes.

## Results

### Checkpoint adaptation drives resistance to genotoxic challenge

To better characterise the cellular consequences of checkpoint adaptation, we established a model where adapted cells could be recovered in a reproducible manner for further analysis. To this end, a previous study has demonstrated that diploid repair-deficient *rad52* mutants readily undergo checkpoint adaptation following exposure to X-rays (Galgoczy and Toczyski, 2001). We performed similar experiments by treating repair-defective homozygous diploids, deleted for the *RAD52* gene, with camptothecin (CPT), which induces double-strand breaks (DSBs) by trapping Top1 cleavage complexes (Top1cc) on the DNA (Hsiang et al., 1989). As expected, in the presence of CPT, *rad52* cells were unable to form colonies on agar plates after 2 days, unlike wild type cells or unchallenged *rad52* cells (Fig. 1a and data not shown). Following 5 days of incubation, small colonies began to form, which were clearly visible after 8 days (Fig. 1a). Despite their growth, the colonies contained a large fraction of metabolically inactive cells as indicated by the accumulation of the vital dye Phloxine B (pink colouring) (Minois et al., 2005). Colony formation was not a result of decreased CPT activity over time, as CPT-containing plates incubated at 30°C for 7 days were as effective in killing *rad52* cells as freshly prepared plates (Supplementary Fig. 1a). Quantification of cell survival normalized to “no drug” control plates revealed that approximately 85 % of the plated *rad52* mutants were able to eventually form colonies following prolonged incubation on CPT solid agar plates (Fig. 1b). To demonstrate that checkpoint adaptation was responsible for the growth of repair-defective *rad52* cells on CPT, we introduced the adaptation-defective *cdc5-ad* allele (Toczyski et al., 1997). Survival of adaptation-deficient *rad52 cdc5-ad* mutants was strongly reduced (to approximately 30 %, Fig. 1a, b), suggesting that checkpoint adaptation is required for the delayed colony formation. In cells that were repair competent, the presence of the *cdc5-ad* allele had no effect on viability following CPT challenge (Fig. 1a, b). Similar results were obtained when X-rays were used as a source of DNA damage (Fig. 1c, Supplementary Fig. 1b) as previously described (Galgoczy and Toczyski, 2001). In response to the radiomimetic drug bleomycin, adaptation-deficient *rad52 cdc5-ad* cells were also significantly sensitized to the drug treatment compared to *rad52 CDC5* mutants (Supplementary Fig. 1c). Although not as penetrant as the *cdc5-ad* allele, mutations in casein kinase II also result in defective adaptation (Toczyski et al., 1997). Accordingly, the deletion of *CKB2,* also resulted in defective colony formation of *rad52* mutants following genotoxic challenge (Supplementary Fig. 1d, e). In conclusion, checkpoint adaptation promotes the delayed survival and growth of repair-defective cells following genotoxic stress.

**Figure 1.**
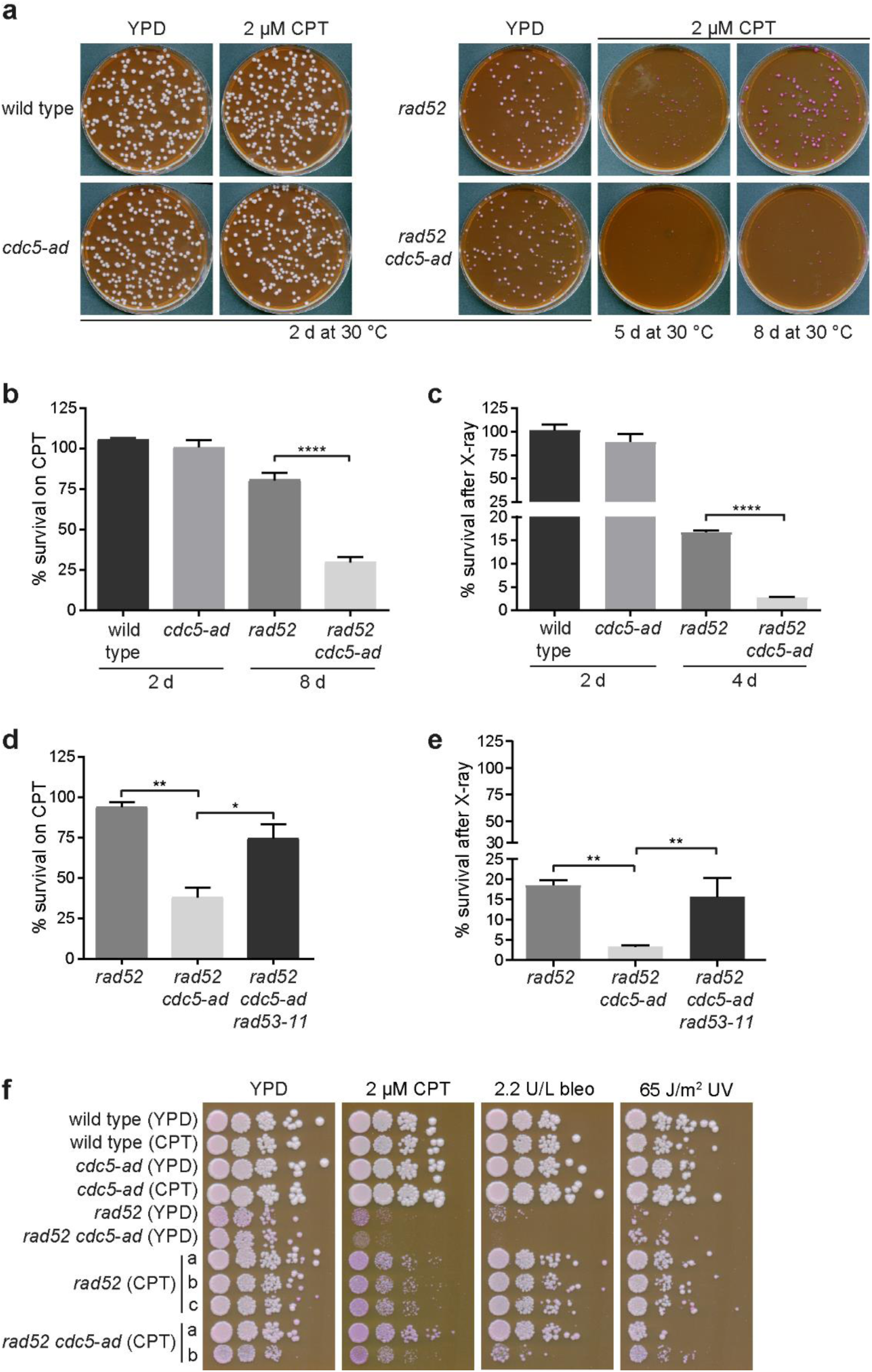
Survival of repair-defective cells after genotoxic stress is facilitated by checkpoint adaptation. (**a**) Representative images of a plating assay. Homozygous diploids were plated onto control agar plates (YPD) or plates containing 2 µM camptothecin (CPT) and incubated at 30 °C for the indicated times. Plates contained 8 µg/mL Phloxine B to visualize cellular metabolic activity. d, days. (**b**) Quantification of colony numbers after plating as described in (a). All visible colonies formed were counted and numbers of colonies formed on CPT-containing plates were normalized to the numbers obtained from YPD plates. Data are represented as mean values and SEM of 3 independent experiments. Statistical analysis was performed using one-way ANOVA with Tukey’s post-hoc test (**** is p ≤ 0.0001). (**c**) Survival of the indicated yeast strains in response to 45 Gy of X-rays was quantified as described in (b). Of note, the cell number plated on X-ray treated plates was ten-times higher than the cell number plated on control plates or to assess cell survival in response to CPT. (**d**) and (**e**) Quantification of cell survival after CPT exposure (d) or in response to X-ray irradiation (e) as described in (b) (* is p ≤ 0.05, ** is p ≤ 0.01). (**f**) Yeast strains of the indicated genotypes derived from either control plates (YPD) or CPT-containing agar plates (CPT) were patched and following over-night growth in liquid YPD, spotted in serial dilutions on agar plates (YPD as a control). Plates contained the indicated genotoxic agents (2 µM CPT or 2.2 units/ L bleomycin (bleo)) or received 65 J/m^2^ UV. Colony formation was monitored after 3 days at 30°C. a-c indicate biological replicates.

To confirm that the *cdc5-ad* associated decreased viability was a result of checkpoint maintenance, we employed the checkpoint-defective *rad53-11* allele (Weinert et al., 1994) to see if it would abolish the enhanced sensitivity phenotype. Indeed, preventing adaptation in *rad52 cdc5-ad* mutants was only able to decrease viability following CPT or X-ray treatment in a checkpoint-competent *RAD53* background but not in *rad53-11* cells (Fig. 1d, e).

To investigate if adapted *rad52* mutants that were able to eventually survive genotoxic stress had acquired drug resistance, we exposed the adapted cells to a second genotoxic challenge. As shown in Fig. 1f, adapted repair-defective mutants derived from CPT-containing plates (*rad52* (CPT)) displayed a significant growth advantage as compared to *rad52* cells that had not been previously exposed to CPT (*rad52* (YPD)). The few *rad52 cdc5-ad* cells that had adapted also acquired drug resistance. Similar drug resistance was observed in adapted cells that had been exposed to X-rays, however, these cells remained sensitive to CPT (Supplementary Fig. 1f). Together, these results suggest that the development of genotoxin resistance in repair-defective cells is promoted by checkpoint adaptation.

### Rapamycin can prevent checkpoint adaptation in repair defective cells

We have previously demonstrated that treating cells with the TORC1 inhibitor rapamycin, prevents checkpoint adaptation in response to DNA damage arising from chronic dysfunctional telomeres (Klermund et al., 2014). To determine whether rapamycin was also able to prevent checkpoint adaptation in diploid *rad52* cells after genotoxic treatment, we analysed cell division in a microcolony assay. G1 or S phase cells were micromanipulated onto agar plates containing the indicated drugs and checkpoint adaptation was scored as the appearance of microcolonies consisting of more than 2 or 4 cell bodies (Fig. 2a) (Dotiwala et al., 2013; Toczyski et al., 1997). The presence of rapamycin significantly reduced the frequency of checkpoint adaptation in response to both X-ray (Fig. 2b) and CPT treatment (Fig. 2c) in repair-defective *rad52* mutants. Even in the absence of exogenous damage (first bar Fig. 2b, c) rapamycin slightly reduced the formation of microcolonies, presumably due to irreparable endogenous damage occurring in *rad52* mutants.

**Figure 2.**
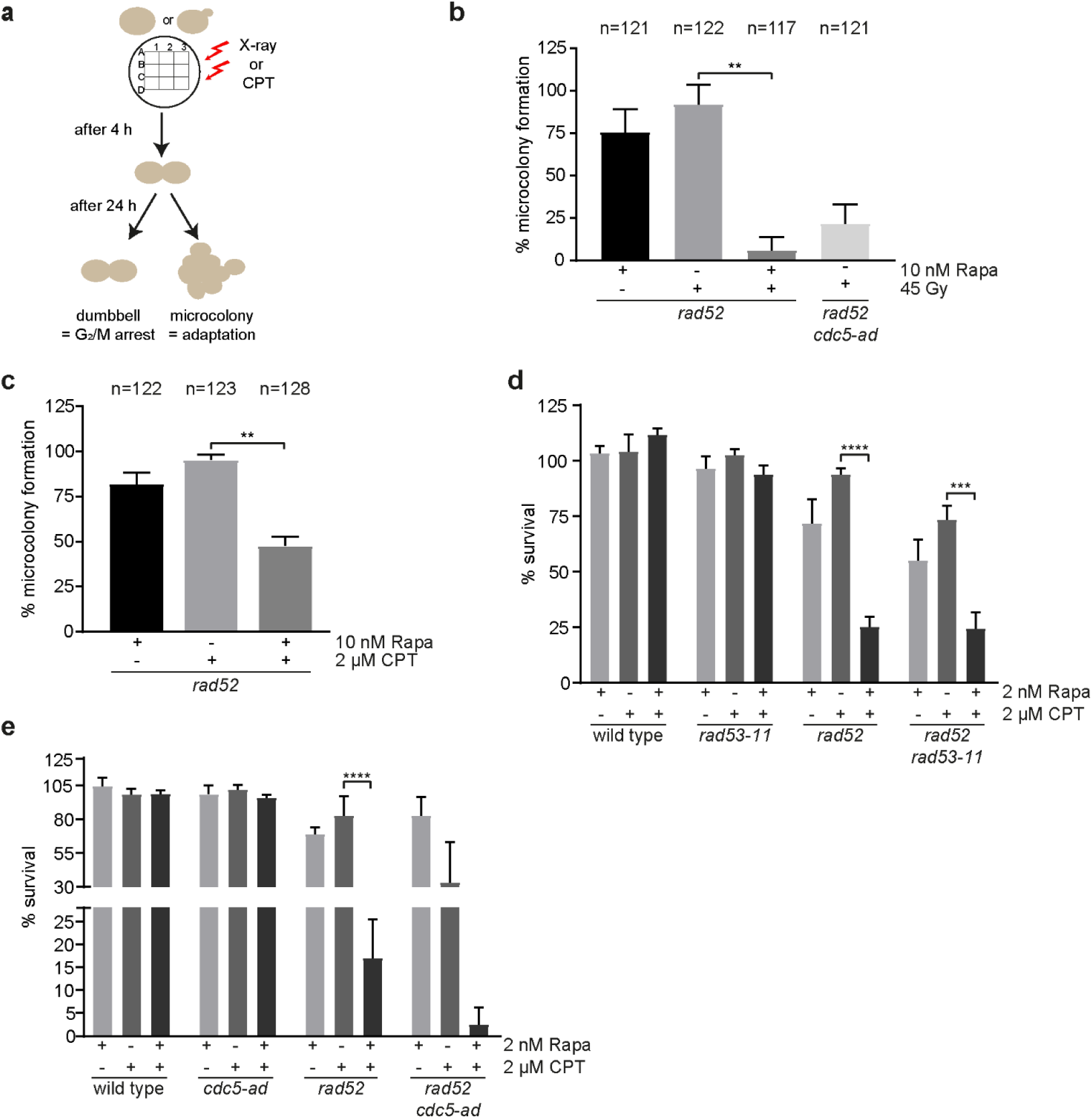
Rapamycin prevents checkpoint adaptation. (**a**) Schematic of the adaptation assay. Following genotoxic challenge (X-ray or CPT), *rad52* mutant cells were scored for their ability to form microcolonies, i.e. undergo checkpoint adaptation (see methods for a detailed explanation). (**b**) As outlined in (a), cells of the indicated genotypes were scored for their ability to adapt in the presence of 10 nM rapamycin or DMSO after irradiation with 45 Gy. Data are represented as mean values and SEM of 5 independent experiments. Statistical analysis was performed using Kruskal-Wallis test followed by Dunn’s post-hoc test (** is p ≤ 0.01). (**c**) As described in (b), microcolony formation was assessed in response to 2 µM CPT in the presence of 10 nM rapamycin or DMSO. Data are represented as mean values with the SEM of 5 independent experiments. (**d, e**) Quantification of cell survival in the presence of the indicated drugs was performed as described in Fig. 1b. Data are represented as mean values with the SEM of 3 independent experiments. Statistical analysis was performed using two-way ANOVA with Tukey’s post-hoc test (*** is p ≤ 0.001 and **** is p ≤ 0.0001).

To test the effects on cell survival, we concomitantly exposed *rad52* cells to rapamycin and CPT, which resulted in approximately 25% cell survival of *rad52* mutants compared to approximately 94 % survival after CPT treatment alone (Fig. 2d). In contrast to the *cdc5-ad* allele (Fig. 1d, e), the decrease in cell survival due to rapamycin was not reversed when the checkpoint was compromised by introducing the *rad53-11* allele (Fig. 2d). Moreover, the presence of rapamycin led to an additive effect in decreasing cell survival of adaptation-deficient *rad52 cdc5-ad* mutants (Fig. 2e). Similar results were obtained when using X-rays as a source of DNA damage instead of CPT (Supplementary Fig. 2a, b).

In summary, rapamycin, like the *cdc5-ad* allele, is able to prevent checkpoint adaptation in repair-defective cells exposed to genotoxic agents. Furthermore, the strong loss of viability when combining either rapamycin (Fig. 2d, e, Supplementary Fig. 2a, b) or *cdc5-ad* (Fig. 1b, c) with genotoxic agents is specific to repair-defective cells, as repair-proficient cells remain fully viable. Unlike the *cdc5-ad* allele, rapamycin is also able to impart toxicity in a checkpoint-independent manner (Fig. 2d, Supplementary Fig. 2a), which can account for the additive cytotoxicity when *rad52 cdc5-ad* cells are exposed to rapamycin in the presence of genotoxins (Fig. 2e, Supplementary Fig. 2b).

### Adapted repair-defective cells undergo extensive chromosome loss

In the absence of *RAD52*, chromosome loss is frequently observed and this is exacerbated upon X-ray treatment (Mortimer et al., 1981). Consistently, DNA content analysis of adapted *rad52* mutants after genotoxic treatment revealed a high degree of chromosome loss resulting in aneuploid cells ranging from near diploid to near haploid DNA contents. This was highly reproducible in the over 50 adapted clones that were tested for DNA content (Fig. 3a and data not shown). Pulsed-field gel electrophoresis indicated that in our conditions, gross chromosomal rearrangements were not occurring in the adapted cells (Supplementary Fig. 3a). By employing a whole-genome sequencing approach, we were able to conclude that the aneuploidy was a result of whole chromosome loss events (Fig. 3b, see Supplementary Fig. 3b for corresponding DNA content FACS). Following CPT treatment of *rad52* cells we observed the loss of one copy of nearly all chromosomes, with the exception of chromosome III (Fig. 3b). To rule out that the reduction in chromosome numbers may have occurred through the induction of meiosis, we deleted the *SPO11* gene in *rad52* mutants (Keeney, 2001), and exposed them to CPT. In the meiosis-defective cells, we observed that the number of adapted cells was not altered (Supplementary Fig. 3c), and moreover, the aneuploid karyotype remained prevalent (Supplementary Fig. 3d).

**Figure 3.**
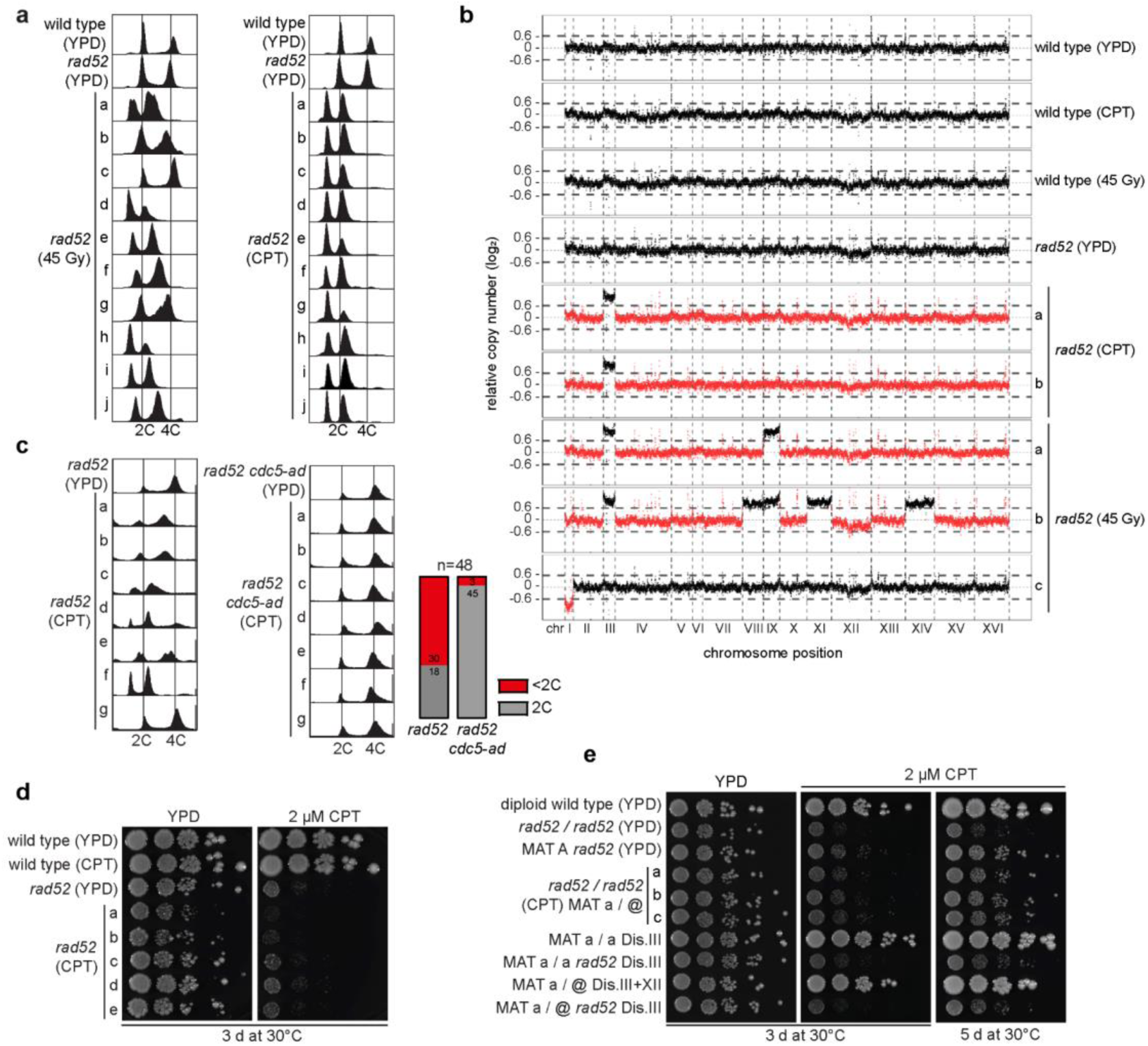
Extensive chromosome loss renders adapted cells aneuploid. (**a**) DNA content of the indicated yeast strains was assessed by flow cytometry. After colonies were formed on the indicated agar plates (YPD as control, 2 µM CPT or 45 Gy X-ray as genotoxic stress), single colonies were grown in liquid media and analysed. (**b**) Representative segmentation plots depicting chromosome copy number variation in *rad52* mutants derived from either control plates (YPD) or after genotoxic treatment (2 µM CPT or 45 Gy, respectively). The segmentation plot depicts the log_2_ ratio of binned reads (reads per million (RPM), y-axis) for the respective chromosome position (x-axis) normalized to the sample median number of reads per bin (RPM). Dashed horizontal lines indicate a threshold of 0.6 or -0.6, respectively, to allow the identification of chromosome loss events which are indicated in red. For more details about normalization, see methods section. (**c**) DNA content of the indicated yeast strains was assessed by flow cytometry. Strains were grown to exponential cultures in YPD media, and treated with DMSO (control) or 20 µM CPT for 4 h. Cells were plated onto YPD plates to recover single colonies, which were grown in liquid media and analysed. Accompanying histogram summarizes the DNA content from the clones analysed. (**d**) Yeast strains of the indicated genotypes from (c) were grown in liquid YPD over-night and spotted in serial dilutions on agar plates (YPD as a control) or plates containing 2 µM CPT. Colony formation was monitored after 3 and 6 days at 30°C. (**e**) Yeast strains of the indicated genotypes were grown in liquid YPD over-night and spotted in serial dilutions on agar plates (YPD as a control) or plates containing 2 µM CPT. Colony formation was monitored after 3 and 5 days at 30°C.

We wondered whether the difference in chromosome loss patterns between *rad52* mutants after X-Ray irradiation and CPT treatment arises due to the different duration of the treatment (acute vs chronic, respectively). Indeed, treatment of *rad52* mutants with a 20 µM CPT dosage for 4 hours yielded a high degree of chromosome loss in 30 out of 48 clones tested, similar to X-Ray exposure (Fig. 3c). Interestingly, the frequency of chromosome loss following acute CPT treatment in adaptation-defective *rad52 cdc5-ad* mutants was drastically lower, with only 3 out of 48 clones showing detectable chromosome loss events (Fig, 3c). In contrast to chronic CPT exposure, that resulted in the formation of CPT resistant *rad52* colonies (Fig.1f), the acute CPT treatment did not lead to resistance (Fig. 3d), suggesting that resistance may be related to the near haploid state and not the aneuploidy, *per se*.

To test this we deleted *RAD52* in strains disomic at chromosome III that were generated by sporulation of triploids (Mulla et al., 2017; Pavelka et al., 2010) and spotted them onto CPT containing plates. Haploid *rad52* strains were more resistant to CPT than diploid mutants, and grew in a similar manner to CPT adapted cells (Figure 3e, first 6 rows). The *rad52* mutants disomic for chromosome III grew similarly to the *rad52* haploids, indicating that aneuploidy did not provide an advantage and is likely not responsible for the resistance. The effects of chromosome III disomy are independent of the mating cassette present (Fig. 3e).

Although we excluded that an extra copy of chromosome III is responsible for genotoxic resistance, we wondered why it was frequently retained in two copies while all other chromosomes were lost. We hypothesized the resistant colonies that arise are heterogeneous in nature, with respect to which chromosomes are lost, however the presence of an extra copy of chromosome III may be advantageous by preventing mating events between aneuploid cells, thereby preventing lethality. Indeed, by isolating single cells (which form single colonies) from the resistant colonies we observed that chromosome III was indeed frequently lost, based on a mating assay (Supplementary Fig. 3f).

This near haploid DNA content of resistant CPT treated cells was stably maintained over several rounds of passaging in liquid media (Supplementary Fig. 3h) and cells did not re-diploidize. In contrast, the X-ray resistant *rad52* mutants underwent progressive chromosome loss and became nearly haploid after several rounds of passaging (Supplementary Fig. 3h). To summarize, following the checkpoint adaptation, the resistant cells that arise undergo extensive chromosome loss which might facilitate their resistance to further genotoxic insults.

### Checkpoint adapted strains become aneuploid and can be targeted pharmacologically

The phenotypes associated with aneuploidy in yeast have largely been described based on the experimental creation of disomic strains (Bonney et al., 2015; Oromendia et al., 2012; Sheltzer et al., 2011; Torres et al., 2007). We tested whether checkpoint adapted aneuploids display similar phenotypes. Due to imbalances in protein complexes disomic strains are defective in maintaining proteostasis, which can be visualized by the inability to resolve Hsp104-GFP positive protein aggregates following heat challenge (Oromendia et al., 2012). Indeed, checkpoint adapted aneuploids (derived from X-ray irradiation) were impaired in resolving Hsp104-GFP foci following 3 hours at 37 °C, when compared to naïve *rad52* mutants and wild type cells (Fig. 4a, b). Consistently, many of these X-ray adapted aneuploids were severely impaired for growth at 37 °C, which correlated with their ability to resolve Hsp104-GFP foci (Fig. 4c). The extent of chromosome loss also served as an indicator for temperature sensitivity (Fig. 4d). The cells that adapted from CPT treatment, which were almost exclusively disomic for chromosome III, but otherwise haploid, did not show defects in Hsp104-GFP foci resolution, nor were they impaired for growth at 37 °C (Supplementary Fig. 4a, b, c).

**Figure 4.**
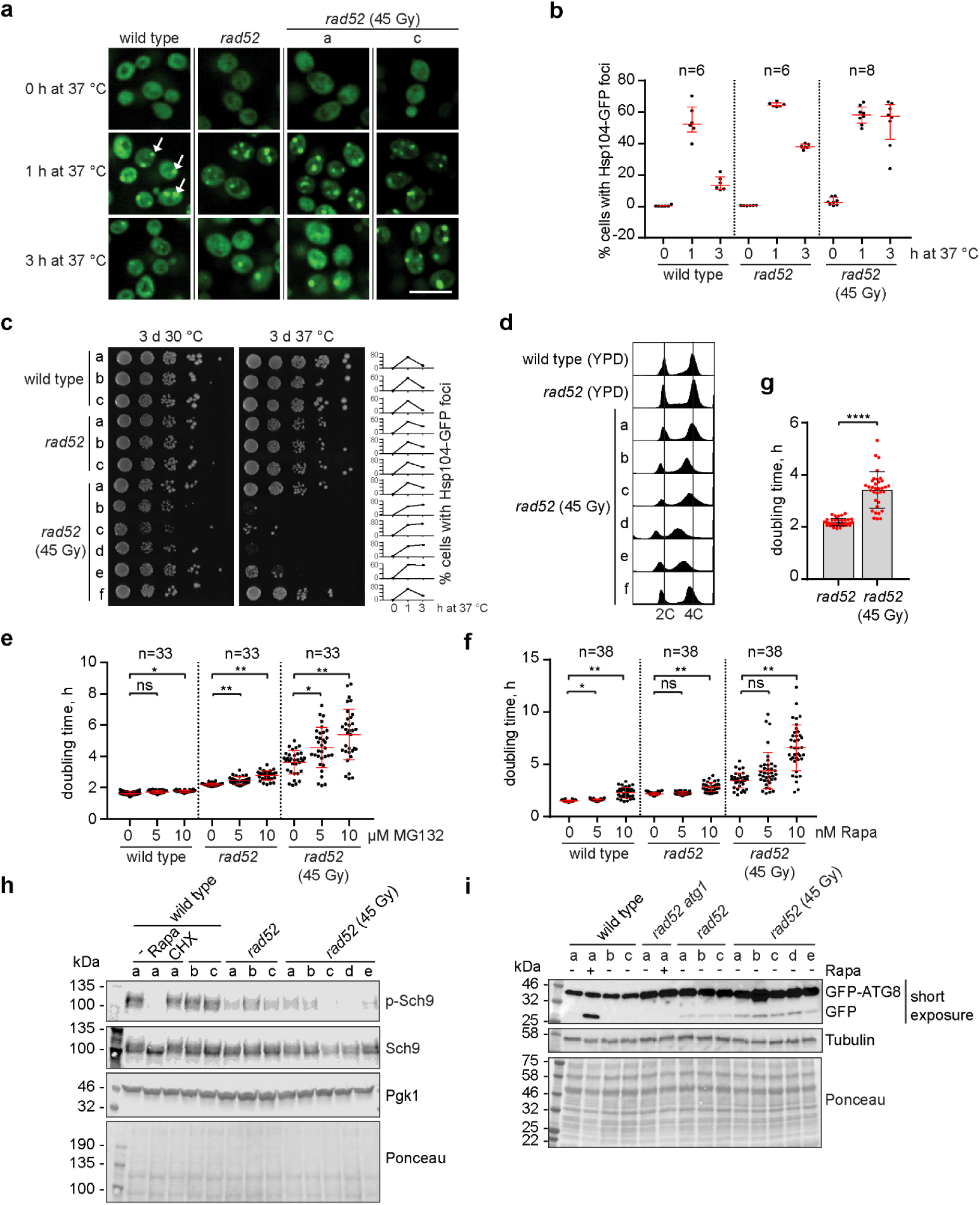
Characterization of aneuploid *rad52* mutants. (**a**) Representative microscopic images of indicated strains upon heat challenge. Control strains and resistant *rad52* colonies were grown to exponential cultures in YPD at 25 °C and exposed to 37 °C for 3 h. All strains expressed endogenous Hsp104 fused with GFP. Arrows indicate Hsp104-GFP foci, for more details on automated foci analysis see methods section. Scale bar, 10 µM. (**b**) Formation of Hsp104-GFP foci in strains from (a) was monitored using fluorescent microscopy. At least 3000 cells (each point on the histogram) per biological replicate were imaged. Error bars indicate median with interquartile range. (**c**) Strains of the indicated genotypes from (a) were spotted in 1:10 serial dilutions on YPD plates. The right panel indicates the percentage of Hsp104-GFP foci upon shift to 37 °C, summarized in (b). (**d**) DNA content of the indicated yeast strains from (a) was assessed by flow cytometry. Letters indicate biological replicates, which are the same clones in panels (a) to (d). (**e**) Population doubling times of control and X-ray resistant *rad52* cells (all in *pdr5* background to allow MG132 uptake) was determined in a growth curve assay in YPD media containing DMSO or the indicated concentrations of MG132 at 30 °C. Data are represented as a scatter plot with indicated mean and SD. Thirty three independent colonies derived from two biological replicates were analysed. Statistical analysis was performed using the Scheirer-Ray-Hare test followed by pairwise Bonferroni-corrected Mann-Whitney-U-tests (* is p ≤ 0.05 and ** is p ≤ 0.001). (**f)** Population doubling times of the indicated strains in YPD media containing DMSO or the indicated concentrations of rapamycin was determined. Thirty eight independent colonies derived from three biological replicates were analysed. Data are represented as a scatter plot indicating mean values and SD and statistical analysis was performed as described in (e). (**g**) Population doubling times of control strains shown in (f) during growth in YPD media containing only the vehicle control DMSO. Data are represented as a scatter plot with bars indicating mean and SD. For statistical analysis, Fligner-Killeen-test for heterogeneity of variances was applied (**** p≤0.00001). (**h**) Indicated strains and X-ray resistant *rad52* colonies were grown to exponential cultures in YPD at 30°C and collected at OD_600nm_ 0.6. Control cultures were either mock-treated, or treated for 15 min with 50 nM rapamycin or 25 µg/mL cyclohexamide. Whole protein lysates were separated by SDS-PAGE and membranes were probed with the indicated antibodies. (**i**) Strains expressing plasmid-encoded GFP-Atg8 were grown to exponential cultures in selective media at 30°C and monitored for autophagy induction. Treatment with 50 nM rapamycin for 1 h served as a positive control. Whole protein lysates were separated by SDS-PAGE and membranes were probed with an anti-GFP antibody.

The fact that aneuploid cells display proteotoxic stress opens an opportunity to target proteostasis regulators in order to induce cytotoxicity (Oromendia et al., 2012; Tang et al., 2011; Torres et al., 2010; Torres et al., 2007). We treated adapted aneuploids with low doses of the proteasome inhibitor, MG132, and observed a striking effect on population doubling time as compared to naïve wild type or *rad52* cells (Fig. 4e). Consistent with the fact that adaptation to CPT did not result in defective Hsp104-GFP foci resolution following heat stress (Supplementary Fig. 4a, b), they also did not show enhanced sensitivity to MG132 (Supplementary Fig. 4d). Apart from sensitivity to proteostatic perturbations, aneuploid cells have also been reported to have altered energy demands, hence rendering them sensitive to TORC1 inhibitors, such as rapamycin (Tang et al., 2011; Torres et al., 2007). Indeed, adaptation following X-ray treatment rendered cells hyper-sensitive to low doses of rapamycin (Fig. 4f). Adaptation to CPT did not result in rapamycin sensitivity (Supplementary Fig. 4e). It has been previously reported that aneuploid cells display characteristic but extremely heterogeneous response patterns after stress (Beach et al., 2017), which can be clearly observed in the responses to MG132 (Fig. 4e) and rapamycin (Fig. 4f). We directly compared the deviations between growth rates in unchallenged conditions and could confirm that there is significant difference between adapted cells and naïve *rad52* mutants (Fig. 4g).

To further understand the nature of the rapamycin sensitivity, we monitored TORC1 activity in checkpoint adapted cells by following the levels of phosphorylated Sch9 (p-Sch9), a direct target of TORC1 (Urban et al., 2007). Whereas TORC1 activity was already reduced in naïve *rad52* cells compared to wild type, the checkpoint adapted aneuploids had even further reduced levels of p-Sch9 (Fig. 4h). Moreover, using the GFP-Atg8 reporter (Tang et al., 2011), we observed increased levels of autophagy in X-ray adapted cells (Fig. 4i) but not in CPT adapted cells (Supplementary Fig. 4f), consistent with the respective rapamycin sensitivity.

In summary, these data demonstrate that although adapted cells acquire genotoxin resistance, their aneuploid status renders them sensitive to agents that target proteostasis and energy regulation pathways.

### CPT adapted cells rely on MRX-mediated DNA damage repair

We next set out to address why checkpoint adapted cells become resistant to further genotoxic challenges. In order to test if adapted *rad52* mutants were still checkpoint-proficient following an adaptation event, we challenged them a second time with either CPT (Fig. 5a) or 45 Gy (Supplementary Fig. 5a). To monitor DNA damage checkpoint activation, we followed the phosphorylation-dependent mobility shift of Rad53 (Lopes et al., 2001) as well as the induction of ribonucleotide reductase 3 (Rnr3) expression (Huang et al., 1998). We observed that all adapted *rad52* mutants had a basal activation of the DNA damage checkpoint, which could be seen in terms of *RNR3* expression, but not Rad53 phosphorylation. Adapted cells were still proficient in checkpoint activation following exposure to the respective DNA damaging agent, ruling out the notion that a faulty checkpoint may allow for the resistance phenotype.

**Figure 5.**
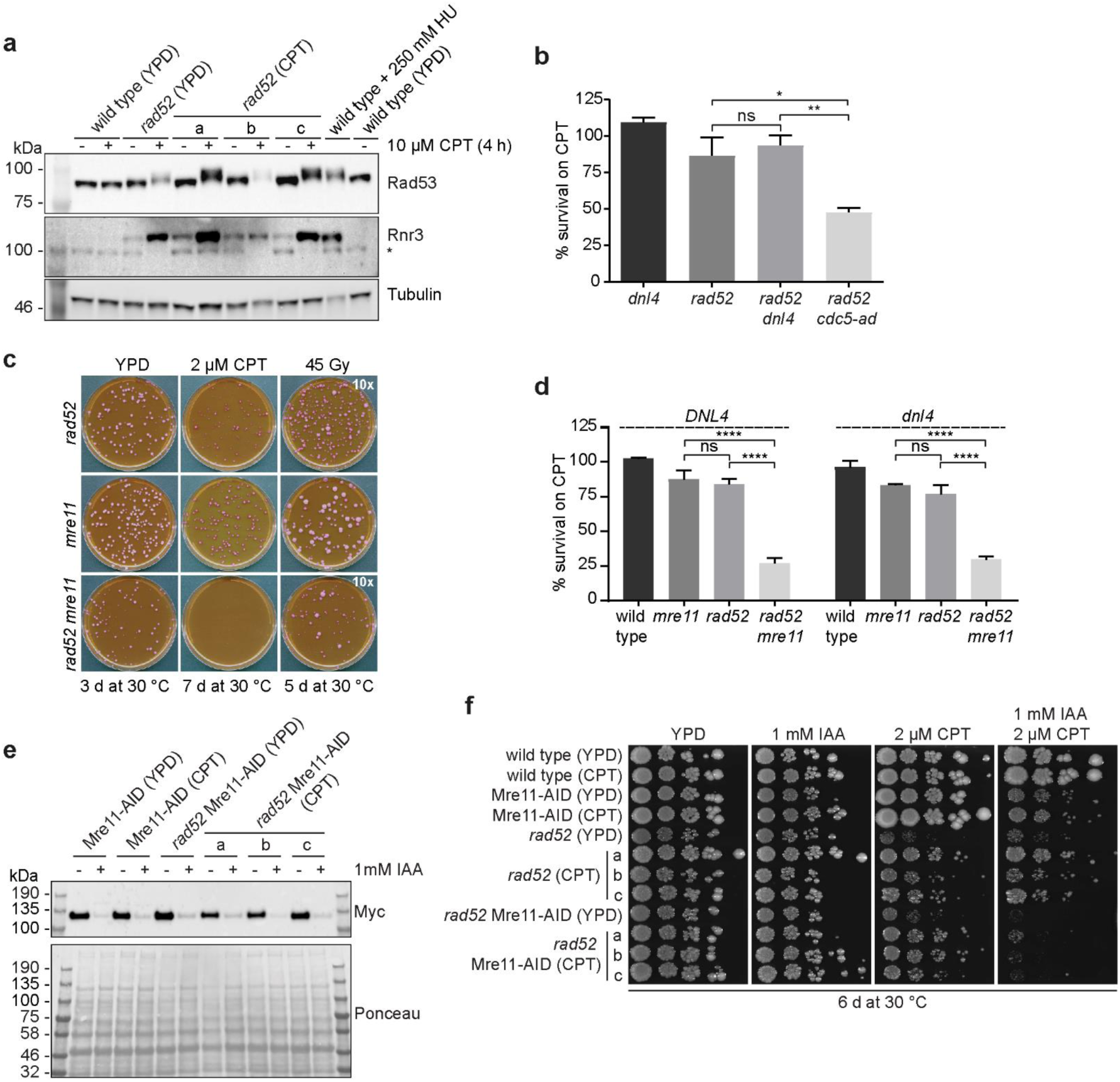
MRX promotes the growth of adapted cells. (**a**) Strains derived from either control plates (YPD) or from CPT treatment (CPT) were grown to exponential phase and treated with 2 µM CPT as indicated. Whole protein lysates were separated by SDS-PAGE and membranes were probed with the indicated antibodies to monitor DNA damage checkpoint activation. Asterisk (*) marks an unspecific band. (**b**) Quantification of cell survival in the presence of 2 µM CPT after normalization to control YPD plates. Data are represented as mean values and SEM of 3 independent experiments. Statistical analysis was performed using one-way ANOVA with Tukey’s post-hoc test (* is p ≤ 0.05 and ** is p ≤ 0.01). (**c**) Representative images of a plating assay. Indicated strains were plated onto YPD or plates containing 2 µM CPT (all containing 8 µg/mL Phloxine B) and incubated at 30°C for the indicated times. Of note, the cell number plated on X-ray treated plates was ten-times higher compared to control plates. (**d**) Quantification of cell survival in the presence of 2 µM CPT after normalization to control YPD plates. The indicated strains were analysed both in a *DNL4*-proficient and *dnl4*-deficient genetic background. Data are represented as mean values and the error bars indicate the SEM of 3 independent experiments. Statistical analysis was performed using one-way ANOVA with Tukey’s post-hoc test (**** is p ≤ 0.0001). (**e**) Strains derived from either control plates (YPD) or plates containing 2 µM CPT were grown to exponential phase and then either mock-treated or treated with 1 mM IAA for 2 h. All strains stably expressed *GDP-AFB2* to facilitate auxin-mediated protein degradation. Whole protein lysates were separated by SDS-PAGE and membranes were probed with the indicated antibodies. (**f**) Strains of the indicated genotype were spotted in serial dilutions and imaged following 6 days of incubation at 30°C.

We therefore tested whether an alternative repair mechanism would allow the survival of post-adaptation *rad52* mutants (Coic et al., 2008; Ma et al., 2003). In addition to homologous recombination, non-homologous end joining (NHEJ) constitutes the second major DSB repair pathway. However, in diploid yeast strains, as used in our experiments, NHEJ is actively suppressed by the transcriptional repression of critical NHEJ factors (Frank-Vaillant and Marcand, 2001; Kegel et al., 2001; Valencia et al., 2001). Nevertheless, since the adapted cells lose chromosomes and resemble a more “haploid-like” karyotype, we sought to address if the survival of *rad52* mutants after genotoxic stress was facilitated by the aberrant use of NHEJ. To test this, we exposed *rad52* mutants lacking DNA ligase IV (Dnl4), a crucial NHEJ factor, to genotoxic stress. The absence of *DNL4* did not compromise cell survival of *rad52* mutants in the presence of CPT (Fig. 5b) or after X-ray irradiation (Supplementary Fig. 5b). We further investigated the involvement of other repair pathways and found that the deletion of either *RAD59, RAD51, RAD55* did not hinder the formation of adapted colonies following either CPT or X-ray exposure in a *rad52* background (Supplementary Fig. 5c-e). Microhomology mediated end joining (MMEJ) is a faulty repair pathway that does not depend on either Rad52 or on the Ku proteins, but rather utilizes the MRX complex, DNA polymerases δ (Pol3, Pol31, and Pol32) and λ (Pol4), and endonuclease Rad1/Rad10 (Ma et al., 2003; Meyer et al., 2015). Strikingly, the deletion of *MRE11,* prevented the growth of adapted cells in the presence of CPT (Fig. 5c, d), and although colony number was not affected colony size was compromised following X-ray exposure (Fig. 5c, Supplementary 5f). Another MRX (Mre11-Rad50-Xrs2) component, Rad50, also contributed to the re-growth of CPT adapted cells (Supplementary Fig. 5g, h). The deletion of *RAD1* and *POL4* did not influence the formation of colonies following CPT or X-Ray treatment (Supplementary Fig. 5i, j). Consistent with these genetic data, we were unable to detect changes in rates of MMEJ using a previously described reporter, when comparing adapted to non-adapted cells (Supplementary Fig. 5k) (Meyer et al., 2015).

Since Mre11 does not contribute to adaptation *per se* (Lee et al., 1998), we considered that it may become essential for post-adaptation repair. To address this, we created an auxin-inducible degron (AID) version of Mre11 (Fig. 5e). We allowed *rad52* cells to adapt and form colonies in the presence of Mre11 and then assessed their sensitivity to CPT following auxin (IAA)-induced removal of Mre11. Indeed, we observed that adapted *rad52* cells acquire resistance to CPT in the presence of Mre11, but become sensitive following the subsequent removal of Mre11 (addition of IAA) (Fig. 5f). The sensitivity exceeds that of losing Mre11 in the presence of Rad52 and is similar to that of non-adapted *rad52* mutants.

Together, these data suggest that the MRX complex contributes to the resistance phenotype of adapted *rad52* cells in response to chronic CPT exposure.

## Discussion

Due to the fact that many human cancers harbour mutations in DNA repair genes, the use of genotoxic agents is frequently effective in the initial reduction of tumor size. Unfortunately, these treatments become ineffective as chemotherapy resistance develops. Therefore, it is essential to characterize the molecular events that occur when repair-defective cells are treated with genotoxins so that the pathways which lead to resistance can also be targeted in a combinatorial manner. We have employed repair-defective diploid yeast (*rad52/rad52)*, as a model to gain insights into these processes. Here, we demonstrate that in response to ionising radiation, as previously shown (Galgoczy and Toczyski, 2001), and CPT treatment, checkpoint adaptation contributes to the eventual re-growth of cells. Importantly, the adapted cells that re-grow are resistant to further genotoxic challenges and undergo extensive whole chromosome losses (Figure 3), translocations and break-induced replication (Galgoczy and Toczyski, 2001). By preventing checkpoint adaptation through expression of *cdc5-ad,* deletion of casein kinase II or rapamycin treatment, the number of resistant colonies that form is drastically reduced. Moreover, rapamycin can also cause cytotoxicity in a post-adaptive manner, due to the aneuploid nature of the cells (Tang et al., 2011; Torres et al., 2007). Taken together, we describe the sequence of events that occur when repair-defective cells acquire drug resistance, and demonstrate options for targeting them either genetically or pharmacologically.

Checkpoint adaptation allows the continued proliferation of cells, despite the presence of chromosomal damage (Sandell and Zakian, 1993; Toczyski et al., 1997). Ideally, repair-competent cells should repair DNA damage before adaptation occurs (recovery), in order to prevent the risks associated with cell proliferation in the presence of lesions (Figure 6). In repair-proficient cells the inhibition of adaptation does not have a negative effect on the sensitivity of the cells to DNA damage (Figure 1), in fact in the case of chronic telomere uncapping (*cdc13-1* cells), preventing adaptation is even advantageous for cell survival as long as the cells are given the opportunity to repair the damage (Klermund et al., 2014). When repair pathways are defective, the only option for cell survival following genotoxic stress is via checkpoint adaptation, therefore the inhibition of checkpoint adaptation in this scenario provides a synthetic lethal-like effect (Figure 6). The extrapolation of these findings to the scenario in human cancer suggests that the combination of adaptation inhibition with genotoxic drugs may give the desired effect of only causing cytotoxicity in the repair-defective tumor cells and not in the repair-proficient healthy tissue. In support of this notion, it has been demonstrated that BRCA1^-/-^ breast cancer cells are hypersensitized to PARP inhibitors (PARPi) in the presence of rapamycin (Osoegawa et al., 2017; Sun et al., 2014), although the role of checkpoint adaptation was not assessed in these studies. Furthermore, inhibition of PLK1 also leads to PARPi hypersensitization in BRCA1^-/-^ cells (Li et al., 2017).

**Figure 6.**
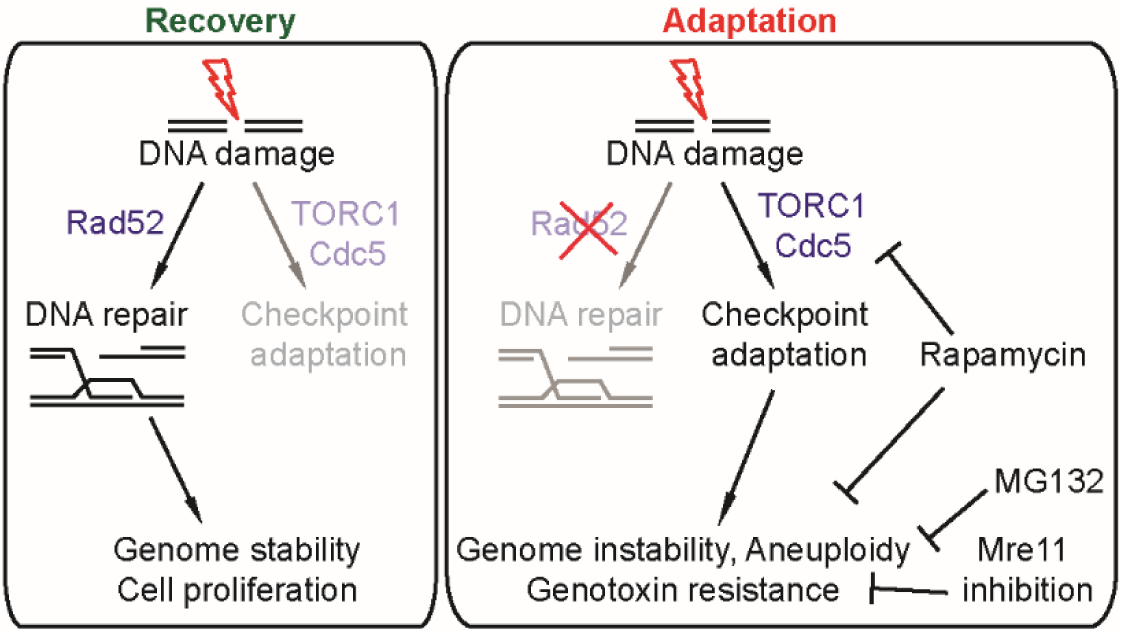
Checkpoint-adapted cells can be targeted. DNA damage activates the DDC to allow time for repair. Rad52 is critical for the repair of DSBs in *S. cerevisiae.* Following successful repair, the checkpoint is turned off and cells can continue to proliferate, referred to as checkpoint recovery (left panel). When damage is not repaired, cells will nonetheless eventually attempt to divide even though the damage persists, by actively extinguishing the checkpoint. This is promoted by TORC1 and Cdc5, both which can be targeted by rapamycin. Following adaptation, cells lose chromosomes and acquire genotoxin resistance. The aneuploid state of these cells renders them sensitive to proteotoxic stress (MG132) and rapamycin. In the case of CPT, their drug resistance can be prevented by disabling the MRX complex.

Aneuploid cells experience proteotoxic stress due to imbalances in multi-subunit protein complexes, rendering them particularly sensitive to perturbations in protein quality control pathways (Donnelly et al., 2014; Oromendia et al., 2012; Tang et al., 2011; Torres et al., 2010; Torres et al., 2007). Consistently we found that aneuploids resulting from adaptation were hypersensitive to the proteasome inhibitor, MG132. Adaptation-derived aneuploid cells were also sensitive to rapamycin (Figure 4), supposedly due to their altered energy demands owing to proteotoxic stress (Stingele et al., 2012; Torres et al., 2007; Williams et al., 2008). Therefore, rapamycin is particularly interesting as it can prevent checkpoint adaptation (Klermund et al., 2014), but can also target aneuploids following adaptation (Figure 6). This explains the observations that in repair-defective cells exposed to genotoxins, rapamycin is additive with the adaptation-defective *cdc5-ad* allele (Fig. 2e) and it still prevents regrowth of checkpoint-defective cells (Fig. 2d). In contrast, the enhanced cytotoxicity of the *cdc5-ad* allele in *rad52* cells depends on the presence of the checkpoint, indicating that its action is solely through the DDC (Fig. 1d, e). Previous studies suggest that rapamycin inhibits Cdc5 function, which is likely how it prevents checkpoint adaptation (Klermund et al., 2014; Nakashima et al., 2008). How rapamycin specifically targets aneuploids remains enigmatic, although we have observed that TORC1 signalling is reduced in adapted cells as compared to naïve *rad52* and wild type cells (Fig. 4d, e).

Interestingly, CPT caused more extensive chromosome loss than X-ray treatment (Fig. 3b) whereby the CPT adapted cells were nearly exclusively disomic, having only one extra chromosome. X-ray adapted cells were very heterogeneous and were more “extreme” in the extent of their chromosome imbalances. These differences were due to the chronic nature of the CPT treatment. Indeed, when CPT was administered in an acute manner, adapted cells displayed karyotypes similar to X-ray treated cells. The degree of chromosome loss was reflected in the extent of the sensitivity of the cells to proteotoxic stress and rapamycin (compare Fig. 3 and 4 with Supplementary Fig. 3 and 4). Consistently, in controlled experimental conditions, it has previously been observed that a greater number of unbalanced chromosomes results in stronger aneuploidy-associated phenotypes (Torres et al., 2007; Torres et al., 2008). It is interesting that chromosome III was consistently retained as the disomic chromosome in the CPT treated cells. We demonstrate that this may have been a selection to prevent the mating of aneuploid cells, which would lead to further genome instability and likely cell death.

Importantly, aneuploidy *per se* does not render cells resistant to DNA damage (Sheltzer et al., 2011)(Fig. 3d, e) and hence we screened for candidate genes that may be responsible for the resistance emerging following adaptation. We observed that the adapted cells rely on the MRX complex for their acquired resistance to CPT, and X-rays. Surprisingly, the resistance phenotype does not rely on compensation from other components of the HR machinery, including *RAD51, RAD55* and *RAD59* (Coic et al., 2008). Therefore, inhibition of Mre11 in combination with CPT provides yet another opportunity to target repair defective cells and prevent the occurrence of aneuploid resistant cells. It has recently been demonstrated that cells undergoing replicative senescence due to telomere shortening also frequently adapt, which leads to subsequent genome instability (Coutelier et al., 2018; Galgoczy and Toczyski, 2001). It is tempting to speculate that by preventing adaptation, it may also lead to the elimination of senescent cell. In agreement, we have previously demonstrated that the longevity extension properties of rapamycin are partially dependent on the DNA damage checkpoint (Klermund et al., 2014).

Taken together, we have used a yeast model to demonstrate the path taken by repair-defective cells in order to achieve drug resistance and by characterizing these cells, have elucidated ways to prevent their occurrence.

## Methods

### Spotting assay

Overnight cultures grown in YPD at 30 °C were spotted in tenfold serial dilutions starting with OD_600nm_ 0.5 in the first dilution onto the indicated agar plates. Images were taken after 3 to 4 days (unless indicated otherwise) using the ChemiDoc™ Touch Imaging System (Bio-Rad). Agar plates contained the vital dye Phloxine B at a final concentration of 8 µg/mL.

### Western Blotting

Protein extraction was performed as described (Klermund et al., 2014). Briefly, cell pellets corresponding to 2 OD_600nm_ units were harvested by centrifugation and resuspended in 150 µL solution 1 (1.85 M NaOH, 1.09 M 2-mercaptoethanol) followed by incubation on ice for 10 min. After the addition of 150 µL of solution 2 (50 % TCA), the suspension was mixed briefly by vortexing and incubated on ice for 10 min. After clearing the samples by centrifugation at 13,000 rpm for 2 min at 4 °C, the pellet was washed in 1 mL acetone, centrifuged at 13,000 rpm for 2 min at 4 °C and resuspended in 100 µL urea buffer (120 mM Tris-HCl pH 6.8, 5 % glycerol, 8 M urea, 143 mM 2-mercaptoethanol, 8 % SDS, bromophenol blue). Resuspension was completed by incubation at 75 °C (or 50 °C when analysing Rad53 phosphorylation) for 5 min. If necessary, 1 M Tris-HCl at pH 7.5 was added to adjust the pH. Prior to loading, samples were centrifuged at 13,000 rpm for 1 min to precipitate insoluble material.

Proteins were separated on Mini-PROTEAN^®^ TGX Stain-Free™ precast gels (Bio-Rad) using 7.5 % polyacrylamide gels for Rad53 detection or a gradient of 4-15 % polyacrylamide for all other proteins in Mini-PROTEAN^®^ electrophoresis chambers (Bio-Rad). Proteins were transferred to 0.45 µm pore size nitrocellulose membranes (GE Healthcare) either by wet blot in transfer buffer (25 mM Tris, 192 mM glycine, 20 % ethanol) using Mini Trans-Blot^®^ Cell chambers (Bio-Rad) or by semi-dry blot using the Trans-Blot^®^ Turbo™ Transfer System (Bio-Rad) with the pre-set 10 min blotting program for high molecular weight proteins. Blocking was performed for at least 1 h at room temperature in 5 % skim milk/PBST (1.37 M NaCl, 0.03 M KCl, 0.08 M Na_2_HPO_4_ x 2 H_2_O, 0.02 M KH_2_PO_4_, pH 7.4 supplemented with 0.1 % Tween-20) followed by incubation with the primary antibodies diluted in the same buffer over night at 4 °C. Membranes were incubated with secondary antibodies for 1 h (HRP-coupled antibodies) or 2 h (fluorescently-labelled antibodies) at room temperature. Proteins were visualized by chemiluminescence using ECL Western Blotting substrate (Pierce) on a ChemiDoc™ Touch imaging system (Bio-Rad). Alternatively, proteins were detected on an Odyssey CLx Imaging system (Licor). All antibodies used are listed in Supplementary table 4.

### DNA content analysis using flow cytometry

Cell pellets equivalent to 0.18 OD_600nm_ units were harvested by centrifugation, washed once in 1 mL water to remove medium and resuspended in 300 µL water. After fixation in 70 % ethanol, cells were centrifuged and washed in 1 mL water. After centrifugation, pellets were resuspended in 500 µL 50 mM Tris-HCl pH 7.5 containing 10 mg/mL RNase A and incubated at 37 °C for 3 h or overnight. After centrifugation, pellets were resuspended in 500 µL 50 mM Tris-HCl pH 7.5 containing 1 mg/mL Proteinase K and incubated at 50 °C for 45 min. Finally, cells were resuspended in 500 µL 50 mM Tris-HCl pH 7.5. Sonification was performed for 10 sec at level 1 using “constant” mode with a 3 mm sonification tip (Branson) or in a Bioruptor Pico (Diagenode) for 2 cycles 30 sec each. Subsequently, 500 µL 50 mM Tris-HCl pH 7.5 containing 2 µM Sytox Green were added (final concentration 1 µM), cells were transferred to FACS tubes (Falcon) and analysed on a FACSVerse flow cytometer (BD) using FACS Suite software (BD).

### Creation of homozygous diploids

Isogenic haploids of mating type a or α, respectively, were transformed using a standard protocol with different plasmids conferring either antibiotic resistance or bearing auxotrophic markers (plasmids are listed in Supplementary Table 3). After mating under standard conditions, homozygous diploid selection was performed using the plasmid-borne markers. For plasmid-free diploid selection, isogenic haploids were mated in liquid YPD medium at 30 °C for 5 h and diploids were picked based on morphology using a MSM 400 dissection microscope (Singer Instruments) followed by confirmation based on DNA content.

### Genotoxic treatment procedures

For X-ray treatment, cells were irradiated with the indicated doses after plating on YPD agar plates using a 0.5 mm aluminium filter to remove low-energy X-rays. The dose rate was kept constant and was set to approximately 0.9 Gy/min at 130 kV and 5 mA. For further analysis, X-ray resistant colonies were picked directly from the irradiated plates. For camptothecin treatment, cells were plated onto YPD agar plates containing the indicated drug concentration. After the formation of resistant colonies, they were patched onto YPD agar plates and afterwards used for further analysis.

### Plating assay and statistical analysis of cell survival

Cells were taken from 5 mL overnight cultures in YPD at 30 °C and cell numbers per mL were assessed by counting in a Neubauer counting chamber. 300 cells were plated in 2 technical replicates and incubated at 30 °C for the indicated times. Plate images were taken with an Epson Perfection V700 scanner. Colonies were counted manually taking all colonies visible by eye into account and the average of the two technical replicates was used for further quantification. Colony numbers on plates with genotoxic treatment were normalized to number of colonies formed on control YPD plates and expressed as % survival. If not indicated otherwise, every quantification represents three biological replicates. Statistical analysis was performed using one-way ANOVA with Tukey’s post-hoc test after confirming normal data distribution (Shapiro-Wilk test) and testing the heterogeneity of variances (Brown-Forsythe test) as built-in analyses in the Prism 7.02 software package (GraphPad). In the case of plating assays where genotype and treatment represented two different factors, normal data distribution and variance heterogeneity were tested as before. Subsequently, a two-way ANOVA was used to confirm statistically significant interaction between the two factors followed by Tukey’s post-hoc test to report statistically significant differences between data sub-groups.

### Microcolony assay

Cells were grown to saturation overnight in YPD medium at 30 °C. For X-rays as a damaging agent, cells were diluted 1:10 in fresh YPD and transferred to a petri dish followed by irradiation with 45 Gy. Afterwards, unbudded or small budded cells were manipulated onto YPD agar plates containing either 10 nM rapamycin (in DMSO) or DMSO. Based on a similar experiment performed in (Toczyski et al., 1997), microcolony formation was assessed by quantifying cell bodies, i.e. a cell arrest in G_2_/M evident as a dumbbell shape was counted as 2 cell bodies. Microcolonies that contained only one cell body at the 4 h time point were excluded from further analysis. After 24 h, cell bodies present in every microcolony were counted. Cells were counted as adapted if the microcolony contained more than 2 cell bodies. Normal data distribution was tested using the Shapiro-Wilk test. Since data were not normally distributed, a Kruskal-Wallis test followed by Dunn’s post-hoc test was employed. In the case of CPT as a damaging agent, cells were counted as adapted if the microcolony contained 3, 5 or more than 5 cell bodies. Four cell bodies were still considered arrested since a fraction of cells did not arrest with only 2 but 4 cell bodies at 4 h. This is due to a CPT-induced arrest in the next cell cycle if cells had already transitioned through S phase at the start of the experiment. Otherwise, quantification was performed in analogy to microcolony formation after X-ray.

### Genomic DNA extraction

Exponentially growing cultures (20 mL of OD_600nm_ 0.7) were harvested by centrifugation, resuspended in 1 mL gDNA buffer 1 (0.9 M sorbitol, 0.1 M EDTA pH 7.5), transferred to a reaction tube and centrifuged at 13,000 rpm for 1 min. After resuspension in 400 µL of gDNA buffer 1 supplemented with 14 mM β-Mercaptoethanol, cell walls were digested by adding 20 µL of lyticase (2.5 mg/mL) for 45 min at 37 °C. Successful creation of spheroblasts was monitored with a light microscope. Subsequently, spheroblasted cells were centrifuged at 13,000 rpm for 1 min and resuspended in 400 µL 1x TE buffer. After addition of 90 µL gDNA buffer 2 (0.27 M EDTA pH 8.5, 0.44 M Tris, 2.2 % SDS), samples were vortexed briefly and incubated at 65 °C for 30 min. Subsequently, 80 µL of 5 M potassium acetate were added and samples were incubated at 4 °C for at least 1 h. Cell residues were eliminated by centrifugation at 14,000 rpm for 15 min at 4 °C and DNA contained in the supernatant was precipitated by addition of 750 µL cold 100 % ethanol. To facilitate precipitation, samples were incubated at −20 °C for 30 min and subsequently centrifuged at 14,000 rpm for 5 min at 4 °C. After discarding the supernatant, DNA contained in the pellet was washed with 1 mL cold 70 % ethanol followed by air-drying the pellet at room temperature. DNA was resuspended gently in 500 µL 1x TE and incubated at 37 °C for 60 min in the presence of 2.5 µL RNase A (10 mg/mL). Subsequently, 500 µL isopropanol were added, the sample was mixed and centrifuged at 14,000 rpm for 15 min at 4 °C. After washing the pellet with 1 mL 70 % ethanol, DNA was air-dried at room temperature and resuspended gently in 50 µL 1x TE. Insoluble material was precipitated and removed following centrifugation at 14,000 rpm for 10 min at 4 °C.

### DNA library preparation and sequencing

1 µg of genomic DNA was brought to a final volume of 50 µL and sheared using a Covaris S2 ultrasonicator system in a microTUBE AFA Fiber 6×16mm, applying the following shearing parameters: duty factor 10 %, intensity 5, 200 cycles per curst, 45 sec, 7 °C water bath temperature. After DNA shearing, a double-size selection was performed using Ampure XP beads with ratios of 0.6:1 (beads:DNA) to exclude larger fragments and 1:1 (beads:DNA) to remove smaller fragments. This procedure enriched the DNA for the 100-500 bp fragments. Purified DNA was quantified using the Qubit dsDNA HS Assay Kit in a Qubit 2.0 fluorometer (Life Technologies) and the DNA size distribution was profiled in a High Sensitivity DNA chip on a 2100 Bioanalyzer (Agilent Technologies).

DNA sequencing library preparation was performed using NuGEŃs Ovation Ultralow System V2 (2014-2017), following the manufacturer’s recommendations. Libraries were prepared with a starting amount of 80 ng of sheared DNA (100-500 bp in size) and were amplified in 8 PCR cycles. Libraries were profiled on a DNA 1000 chip on a 2100 Bioanalyzer (Agilent Technologies) and quantified using the Qubit dsDNA HS Assay Kit in a Qubit 2.0 fluorometer (Life Technologies).

DNA sequencing (DNA-seq) libraries were sequenced on Illumina NextSeq 500 (75-nt paired-end, with an effective read length of 79 nt), yielding 10-13 million read pairs per sample.

### DNA sequencing analysis

Sequencing qualities were checked using FastQC (version 0.11.5) (https://www.bioinformatics.babraham.ac.uk/projects/fastqc/). All reads were mapped to the yeast genome (Ensembl genome assembly version R64) with the Burrows-Wheeler Aligner BWA (version 0.7.15) (Li and Durbin, 2010). Reads not mapped in a proper pair as well as secondary alignments were removed using Samtools (version 1.3.1) (Li et al., 2009). Furthermore, duplicate reads were removed using MarkDuplicates of the Picard tool package (version 2.6.0) (http://broadinstitute.github.io/picard/). Due to very high and unequal coverage, the mitochondrial chromosome was removed from all subsequent analyses. Reads were counted and summarized in 1000 bp bins using readCounter of the HMMcopy package (version 0.1.1) ((Ha et al., 2012) and (https://github.com/shahcompbio/hmmcopy_utils)). Resulting wig files were transformed to bedgraph files using wigToBedGraph of the kentUtils package (version v302) without collapsing adjacent windows (https://github.com/ENCODE-DCC/kentUtils) and normalized to reads per million (PRM). Using a custom made R script, RPM of each bin were then normalized to the sample-wide median RPM and chromosome copy numbers were ^e^stimated for each sample based on the chromosome median of this bin-wise RPM ratio (log_2_). A cut-off of 0.6 or -0.6 was chosen to determine a higher or lower chromosome copy number than the other chromosomes in the same sample.

### Pulsed-field gel electrophoresis

Exponentially growing cultures corresponding to 25 OD_600nm_ units were collected by centrifugation at 5,000 rpm for 5 min and subsequently washed in 20 mL of 10 mM Tris-HCl, 50 mM EDTA, pH 7.5. Cells were resuspended in 500 µL spheroblasting buffer (0.056 M sodium phosphate, 0.2 M EDTA, 40 mM DTT) and preheated to 42 °C on a heat block. 2 % (w/v) low-melting point agarose in H_2_O was boiled and kept at 42 °C, and 500 µL (1:1) were mixed with the cells in spheroblasting buffer. 90-100 µL of the mix were pipetted into each plug mold (gives 10-12 plugs per sample). Once solidified at 4 °C, the plugs of each strain were transferred to large Falcons (50 mL) containing 3 mL of spheroblasting buffer supplemented with 0.08 mg/mL zymolyase. Plugs were incubated for 24 h at 37 °C, shaking. The plugs were washed with 10 mL 1 x TE buffer. Subsequently, 2 mL of Proteinase K buffer (10 mM Tris-HCl, 0.42 M EDTA, 1 % N-lauroyl sarcosine, 2 mg/mL Proteinase K) were added and the plugs were incubated for a further 24 h at 50 °C, shaking. The plugs were washed five times with 10 mL of 10 mM Tris-HCl, 50 mM EDTA, pH 7.5. Plugs were stored in the same buffer at 4 °C until analysis. For the analysis, plugs were loaded on a 1 % agarose gel and sealed with 1 % agarose. Molecular weight marker CHEF DNA Size Marker #1703605 (Bio-Rad) was used. The gel was run in a CHEF-DR® III system using the following settings: initial switch time 60 sec, final switch time 120 sec, run time 24 h, 6 volts/cm, included angle 120°. The gel was stained with ethidium bromide (1.25 µg/mL in TBE) for 30 min and destained for 10 min before imaging on a ChemiDoc™ Touch imaging system (Bio-Rad).

### Hsp104-GFP microscopy

Overnight cultures were grown in YPD media at 25 °C. Cultures were diluted to 0.2 OD_600nm_ units in 5 mL YPD media, and were grown for 3 h at 25 °C and afterwards shifted to 37 °C. One mL of liquid culture was collected per time point by centrifugation at 3,000 rpm for 3 min. ^C^ells were fixed for 10 min at room temperature in Fixation solution (0.027 M K_2_HPO_4_, 0.073 M KH_2_PO_4_, 2.5 % formaldehyde, pH 6.4), washed twice with Potassium phosphate buffer 1 (0.038 M K_2_HPO_4_, 0.062 M KH_2_PO_4_, pH 6.6), and stored in Potassium phosphate buffer 2 (0.08 M K_2_HPO_4_, 0.02 M KH_2_PO_4_, pH 7.4) at 4 °C. Prior to microscopy, cells were permeabilized with 80 % ethanol for 10 min at room temperature, and resuspended in 500 µL Potassium phosphate buffer 2 containing 0.5 µg/mL DAPI (Sigma-Aldrich). Samples were loaded onto Concavanin A (Sigma-Aldrich) pre-coated 96-well plates (CellCarrier®, PerkinElmer).

Imaging was performed using High-Content Screening Opera Phenix^TM^ microscope (PerkinElmer) in confocal mode, using a water objective with 63x magnification. Fifteen independent fields with five planes each were imaged for each sample. Acquired images were analysed with Harmony High-Content Imaging and Analysis Software (version 4.4, PerkinElmer) using standard building blocks. First, nuclei were identified with algorithm M based on the DAPI signal. Second, cytoplasm was identified with algorithm A based on GFP signal. Cellular morphology properties were calculated with standard built-in algorithm, and cell roundness was set to 65 %. Spot detection of GFP foci was performed in the cytoplasm region of the cell with algorithm C. Cells on the periphery of images were excluded from analysis. We analysed data on the single cell level, with at least 3,000 cells per sample. The percentage of cell population that had Hsp104-GFP foci was calculated using Microsoft Excel 2013, and the data were visualized with the Prism 7.02 software package (GraphPad). Representative images were processed using ImageJ 1.52g software.

### Growth curve assay and statistical analysis

Overnight cultures were grown in YPD media at 30 °C, afterwards the OD_600nm_ was measured in 100 µL culture volume in 96 well plate (Tissue culture plate, Falcon®) using Spark® microplate reader (TECAN). The cultures were diluted in 3 technical replicates to 0.05 OD_600nm_ units in 100 µL YPD with addition of either DMSO, or respective drugs. Cultures were grown at 30 °C overnight with orbital shaking in a 96 well plate (Tissue culture plate, Falcon®) in the Spark® microplate reader, and the growth dynamics of the cultures was monitored by measuring OD_600nm_ every hour. The population doubling time was calculated from an average values of technical replicates at the exponential part of the growth curve using an algorithm from (Roth V. 2006), Doubling Time Computing, available from: http://www.doubling-time.com/compute.php

For the statistical analysis, groups were randomly downsampled to the size of the smallest group. The normality of the data was tested using the Shapiro-Wilk test, and to test for homogeneity of variances, the Fligner-Killeen test was applied. Because the data were neither normally distributed nor had equal variances, the Scheirer-Ray-Hare test (an extension of the Kruskal-Wallis test and, thus, a non-parametric alternative to the two-way ANOVA (Scheirer et al., 1976)) was applied and followed by Dunn’s post-hoc test. For all pairwise comparisons between the groups, the Mann-Whitney U test with Bonferroni correction was applied.

### MMEJ reporter assay

We performed MMEJ reporter assay as described (Meyer et al., 2015Meyer et al., 2015) with a minor modifications. Overnight cultures were inoculated in 5 mL YP-Raffinose (1 % yeast extract, 2 % peptone and 2 % raffinose) and grown at 30 °C. In the morning, the media was exchanged to YP-RaffinoseGalactose (1 % yeast extract, 2 % peptone, 1 % raffinose and 2 % galactose) to induce the expression of HO nuclease. After 4 h of induction, cells were resuspended in 2 mL of YP-Raffinose and plated onto SD-His plates to select for histidine prototrophs. The 1:10 000 dilution of cultures were plated on YPD to estimate the total number of viable cells. Plates were incubated at 30 °C for 4 to 6 days. MMEJ rates were calculated by dividing the number of histidine prototrophs by the number of viable cells. The median frequency of interchromosomal MMEJ was estimated based on 24 to 30 independent colonies.

## Acknowledgements

This work was supported through a generous donation from the Chica-Heinz-Schaller foundation. We are grateful to the IMB media-lab Core Facility, the Genomics Core Facility and the use of its NextSeq500 (INST 247/870-1 FUGG), the Microscopy Core Facility, for constant support and assistance of this study, and the Flow Cytometry Core Facility, for the help with experimental set-up. We thank Robbie Loewith for Sch9 and p-Sch9 antibodies, Bernd Bukau for the Hsp104-GFP strain, Wolf-Dietrich Heyer for the strains with MMEJ reporter, and Rong Li for the aneuploid strains. BL is supported through a Heisenberg Professorship from the DFG (LU 1709/2-1). The authors declare that they do not have a conflict of interest.

**Supplementary Figure 1.**
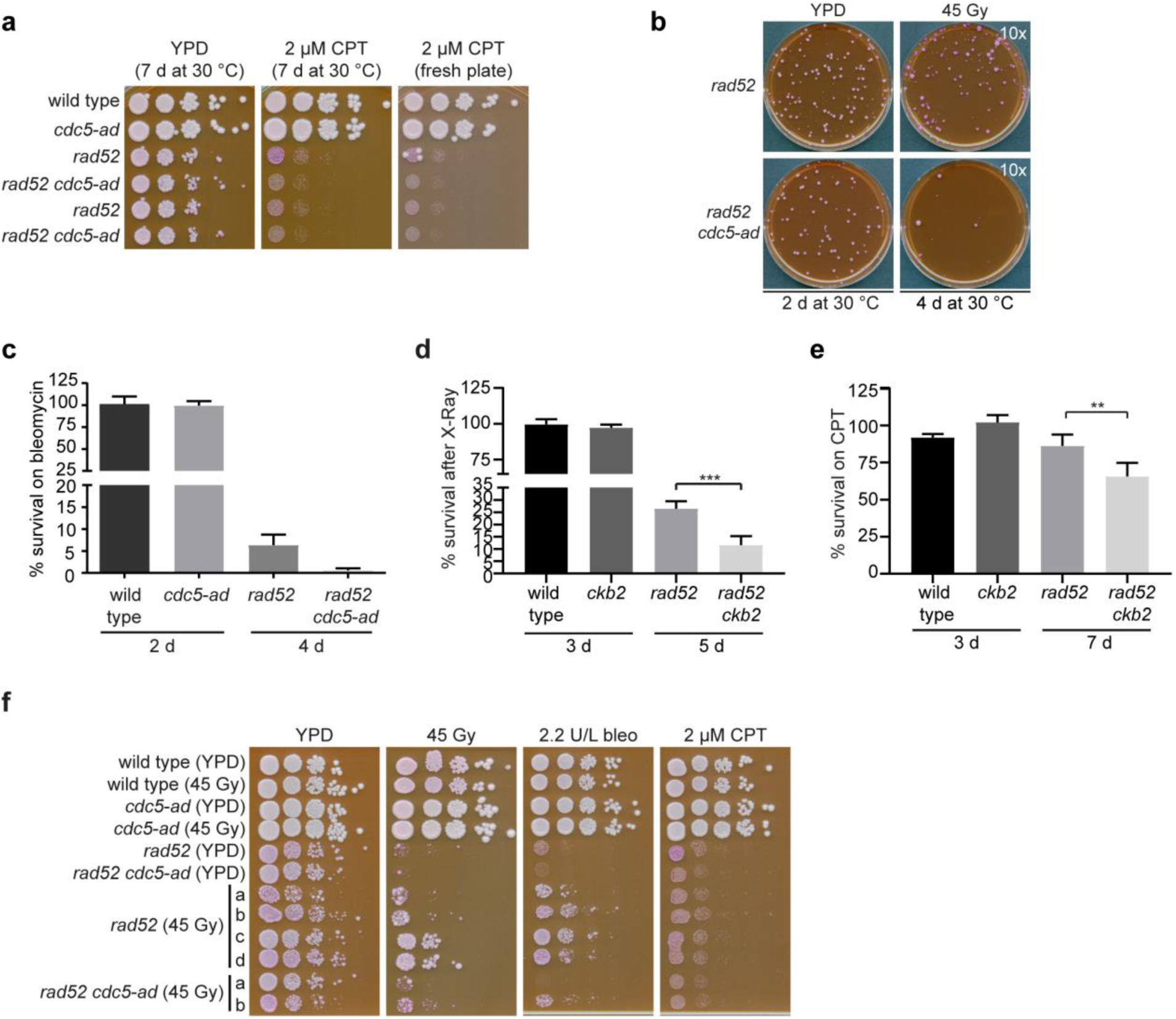
Resistance develops following checkpoint adaptation. **(a)** Yeast strains were spotted in serial dilutions on agar plates. To confirm the continued toxicity of CPT after long-term incubation at 30 °C, cells were spotted onto a CPT-containing plate that had been pre-incubated at 30 °C for 7 days prior to the experiment. A freshly prepared CPT-containing plate and a YPD plate served as controls. (**b**) Representative images of a plating assay. The indicated strains were plated on YPD plates and either left untreated (control) or irradiated with 45 Gy X-rays. Colony numbers were quantified after the indicated incubation times. (**c**) Quantification of cell survival in response to bleomycin treatment. The denoted strains were plated on either control plates (YPD) or plates containing 2.2 U/L bleomycin and colony numbers were assessed and quantified as described previously. Each bar represents the mean value and SEM of 3 independent experiments. (**d, e**) Quantification of colony numbers after plating. (**f**) Analogous to (Fig. 1f), yeast cells derived from either control plates (YPD) or X-ray treated plates (45 Gy) were spotted in serial dilutions onto agar plates. The plates contained either 2.2 U/L bleomycin, 2 µM CPT or were subsequently subjected to 45 Gy of X-ray irradiation. A YPD plate served as a control.

**Supplementary Figure 2.**
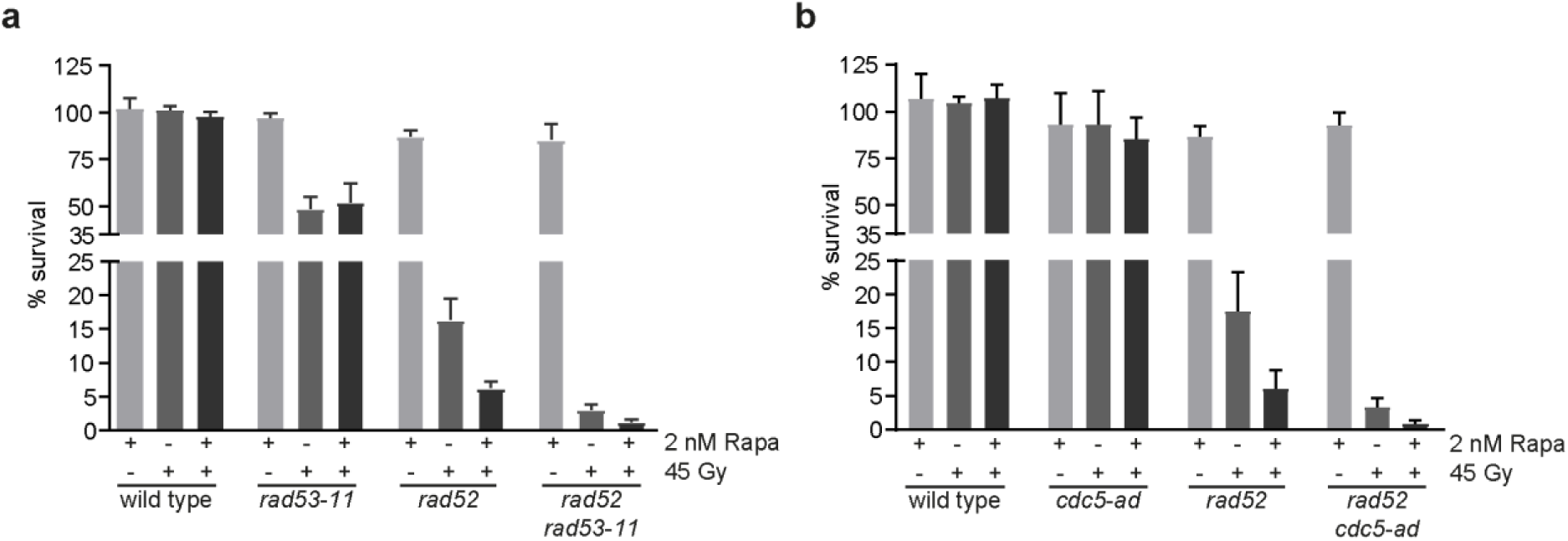
Rapamycin prevents checkpoint adaptation. (**a, b**) The indicated genotypes were scored in an identical manner as described in Figures 2d and e. 45 Gy X-ray exposure was used as a source of genotoxic stress in place of CPT. Each bar represents the mean value and SEM of 4 (a) or 3 (b) independent experiments.

**Supplementary Figure 3.**
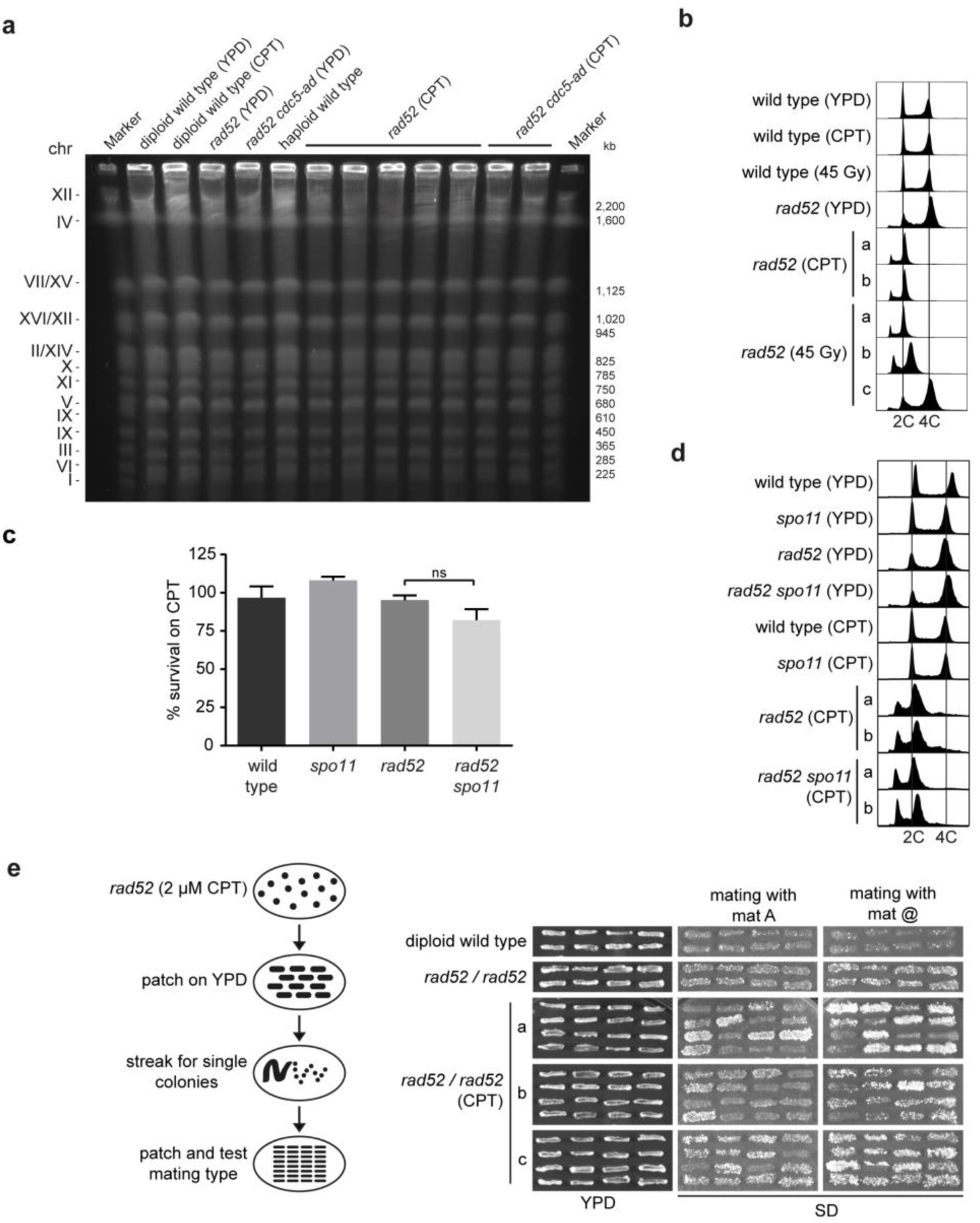
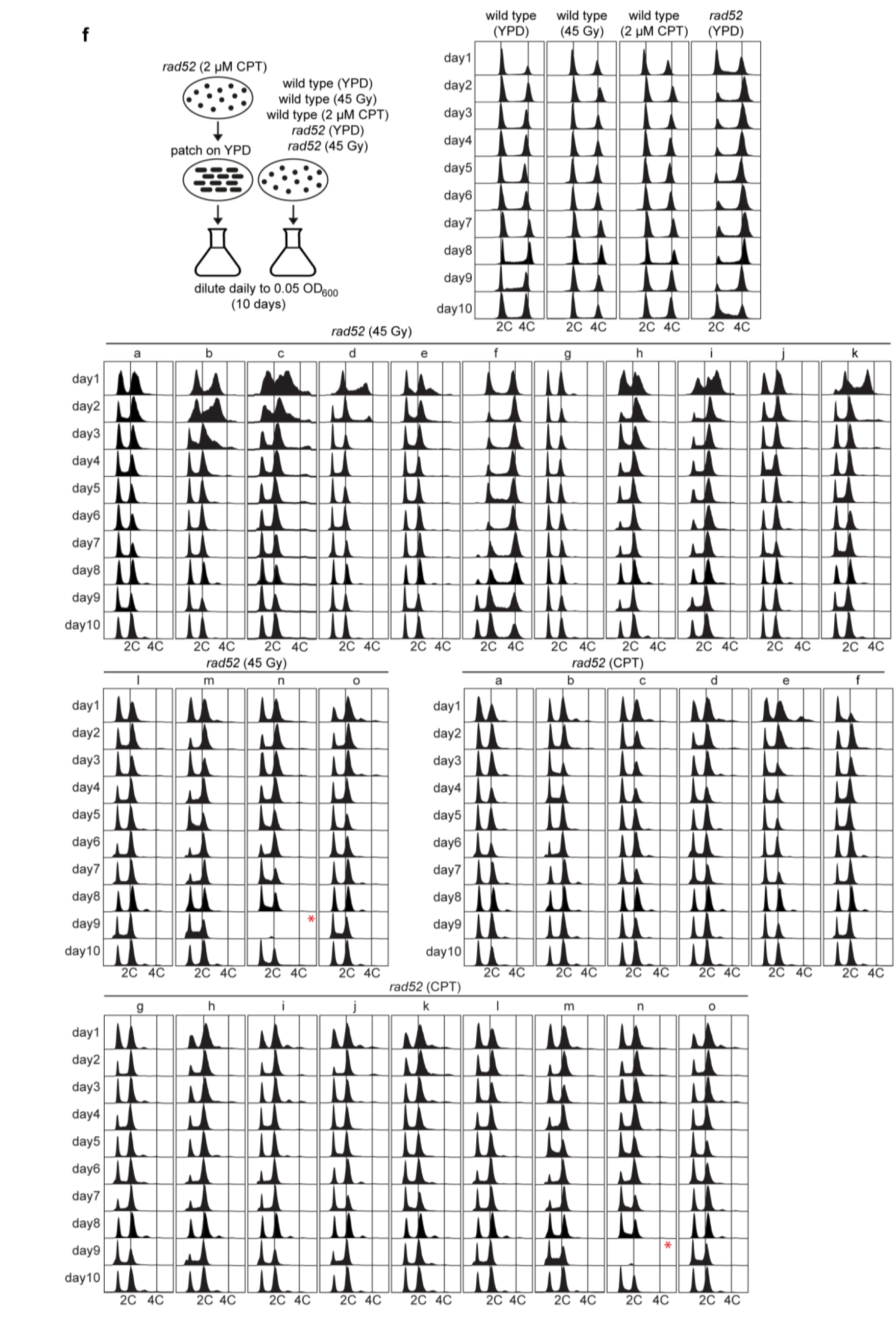
Adapted *rad52* mutants undergo progressive chromosome loss. **(a)** Whole chromosome structure was analysed by pulsed-field gel electrophoresis (PFGE) for the indicated genotypes derived from control conditions (YPD) or after genotoxic stress (2 µM CPT or 45 Gy X-ray, respectively). Roman numerals refer to chromosome number. (**b**) DNA content of the yeast strains corresponding to (Fig. 3b) was assessed by flow cytometry. After colonies were formed on the indicated plates (YPD as control, 2 µM CPT or 45 Gy X-ray as genotoxic stress), single colonies were grown in liquid media and analysed. (**c**) Cell survival in response to 2 µM CPT was assessed in meiosis-deficient cells. rad52 as well as rad52 spo11 mutants and control strains were plated onto CPT-containing or control plates and colony numbers were quantified as described previously. Each bar represents the mean value and SEM of 3 independent experiments. (**d**) Flow cytometry analysis for the adapted cells described in (c). (**e**) Characterization of chromosome III loss frequency in rad52 strains treated with 2 µM CPT by estimating the frequency of mating to a mating tester strains. Ten independent clones of rad52 from plates containing 2 µM CPT were patched on YPD plates to recover the viable cells, streaked for at least 100 single colonies from each patch, and subjected to mating type analysis. Diploid wild type and diploid rad52 from YPD plates served as a control. The strains were mated with mat A and mat @ mating tester strains, and resulting diploids were selected on SD plates. Representative images of mating type analysis are shown. (**f**) Analysis of chromosome loss dynamics in rad52 strains after 2 µM CPT or 45 Gy X-ray exposure. Single colonies of indicated strains were grown over-night in YPD media, and diluted daily to 0.05 O.D_600_. DNA content was assessed by flow cytometry. For the wild type strains derived from YPD plates and plates containing 2 µM CPT or exposed to 45 Gy X-Ray dose, and rad52 strains from YPD plates, five independent colonies were analysed. For the rad52 strains derived from plates containing 2 µM CPT or exposed to 45 Gy X-Ray dose, fifteen independent colonies were analysed. Asterisk (*) indicates missing samples.

**Supplementary Figure 4.**
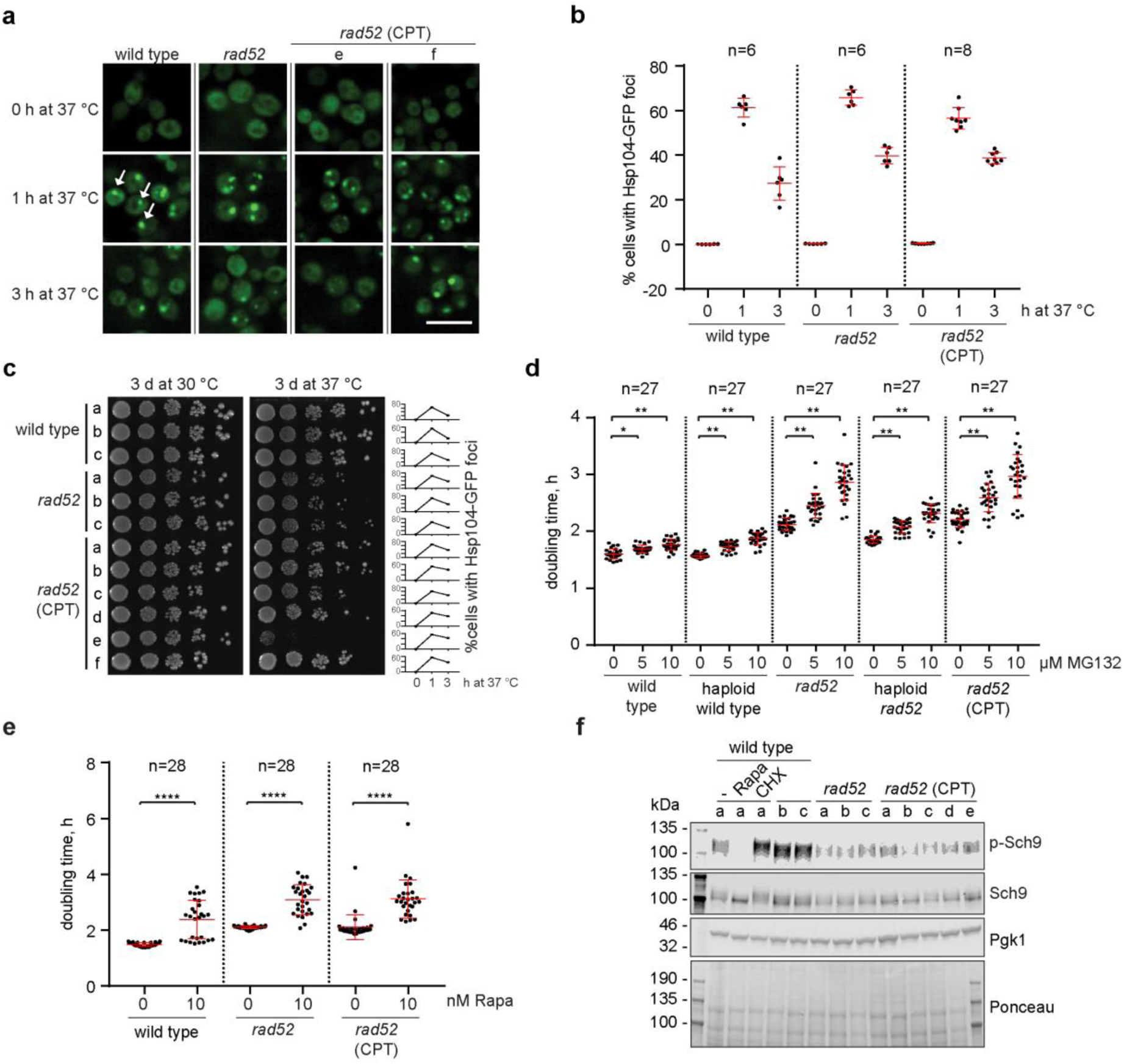
Exploiting aneuploid phenotypes in *rad52* mutants. (**a**) Representative microscopic images of the indicated strains upon heat challenge. All strains expressed endogenous Hsp104 fused with GFP. Arrows indicate Hsp104-GFP foci, for more details on automated foci analysis see methods. Scale bar, 10 µM. (**b**) Six independent colonies of each genotype and eight resistant *rad52* mutants were grown in YPD at 25 °C and exposed to 37 °C. Formation of Hsp104-GFP foci was monitored using fluorescent microscopy. At least 3000 cells per sample (each point on the histogram) were imaged. Error bars indicate median with interquartile range. **(c)** Strains of the indicated genotypes from (b) were spotted in serial dilutions onto YPD plates. The right panel indicates the percentage of Hsp104-GFP foci upon shift to 37 °C, summarized and quantified in (b). Letters indicate single colonies, which are the same clones in panels (a) to (c). (**d**) Growth curve to determine population doubling times of the indicated haploid and diploid strains as well as CPT-resistant rad52 mutants (all in pdr5 background) in YPD media containing DMSO or indicated concentrations of MG132 at 30 °C. Twenty seven independent colonies derived from two biological replicates were analysed. Data are represented as mean and SD in a scatter plot. Statistical analysis was performed using the Scheirer-Ray-Hare test followed by pairwise Bonferroni-corrected post-hoc Mann-Whitney-U-tests (* is p ≤ 0.05 and ** is p ≤ 0.001). (**e**) Population doubling times of the indicated strains and CPT-resistant rad52 cells in YPD media containing DMSO or the indicated concentrations of rapamycin at 30 °C. Twenty eight independent colonies derived from three biological replicates were analysed. Statistical analysis was performed as described in (d) (**** is p ≤ 0.00001). (f) Indicated strains and CPT resistant rad52 colonies were grown to exponential cultures in YPD at 30 °C and collected at OD_600nm_ 0.6. Cultures were either mock-treated or treated for 15 min with 50 nM rapamycin (Rapa) or 25 µg/mL cyclohexamide (CHX). Whole protein lysates were separated by SDS-PAGE and membranes were probed with the indicated antibodies.

**Supplementary Figure 5.**
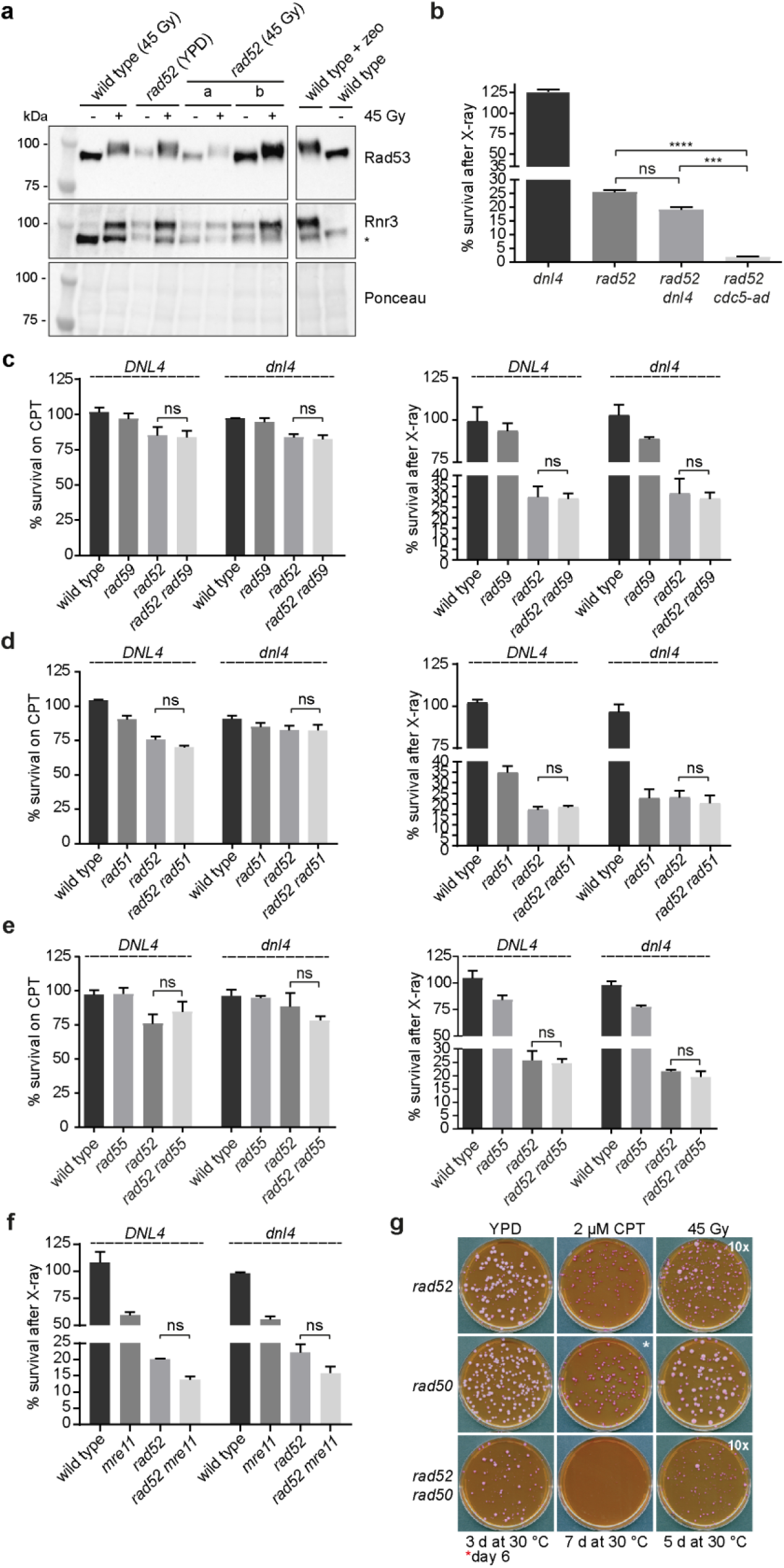
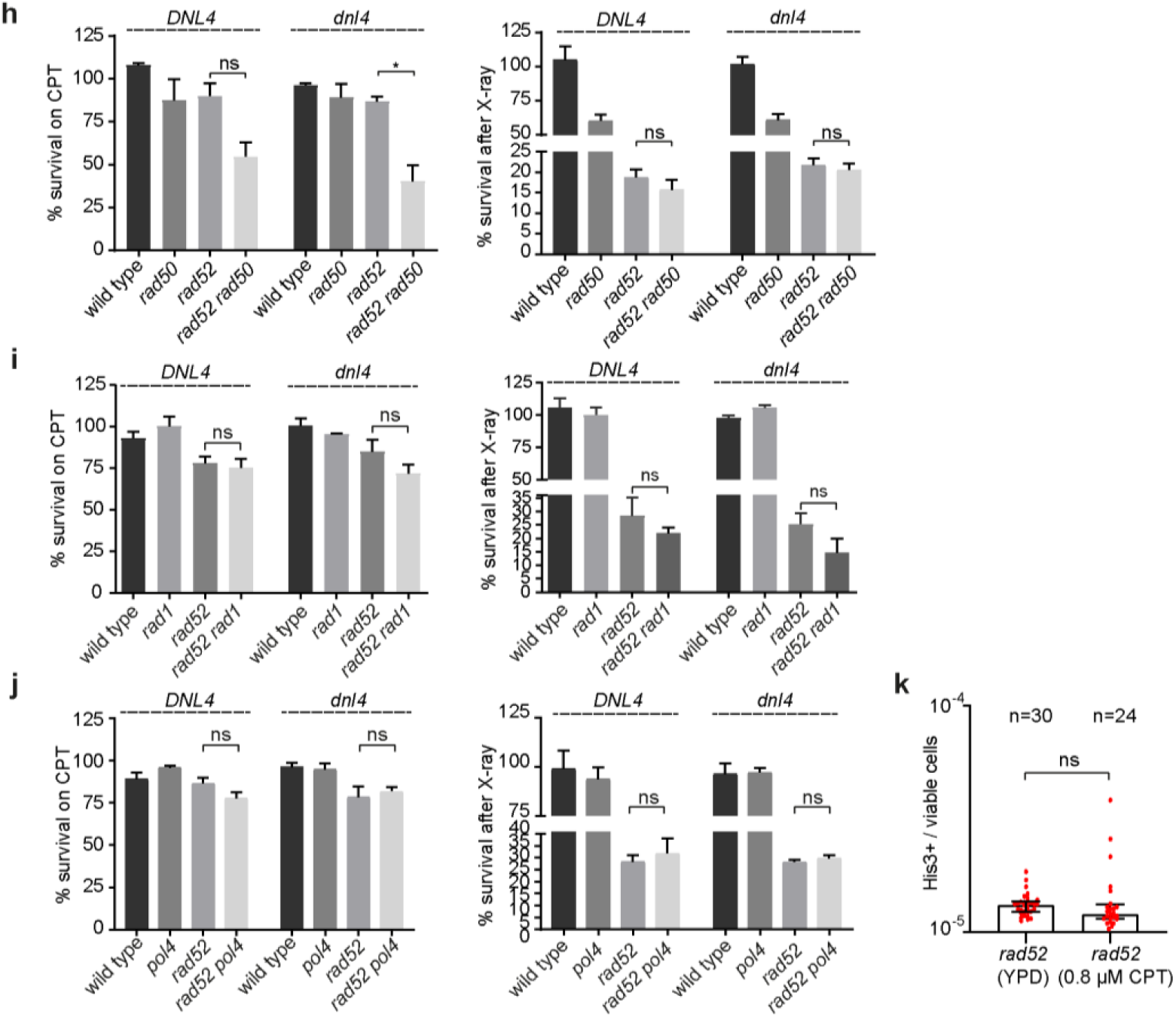
Contribution of repair factors to survival of *rad52* mutants. (**a**) Strains derived from either control plates (YPD) or after 45 Gy of X-ray were grown to exponential phase and then treated with X-rays (45 Gy) as indicated. Whole protein lysates were separated by SDS-PAGE and membranes were probed with the indicated antibodies. Treatment of wild type cells with 250 µg/mL zeocin (zeo) was used as a positive control. (**b**) Analogous to Figure 5b, cell survival in response to 45 Gy of X-ray was quantified as described previously. (**c-f**) Cell survival in response to 2 µM CPT (left panels) or 45 Gy of X-rays (right panels) was quantified as described previously for the indicated strains both in a *DNL4* and *dnl4* background. Data are represented as mean values with the SEM of 3 independent experiments. (**g**) Representative images of a plating assay. Indicated strains were plated onto YPD or plates containing 2 µM CPT (all containing 8 µg/mL Phloxine B) and incubated at 30°C for the indicated times. Of note, the cell number plated on X-ray treated plates was ten-times higher compared to control plates. (**h-j**) Quantification of cell survival in the presence of 2 µM CPT or after exposure to 45 Gy after normalization to control YPD plates. All data are represented as mean values and the error bars indicate the SEM of 3 independent experiments. (**k**) Quantification of interchromosomal MMEJ frequency between his3Δ3’ and his3Δ5’ substrates that share 20 bp microhomology by normalization of cell survival on SD-His to control YPD plates. Strains derived from either control plates (YPD) or plates containing 0.8 µM CPT were grown over-night in YP-Raff. The HO expression was induced by switch to galactose-containing medium for 4 h, and then cells were plated onto SD-His or YPD plates. Data are represented as median values and the error bars indicate interquartile range. Statistical analysis was performed using Student’s t-test.

**Supplementary Table 1.**
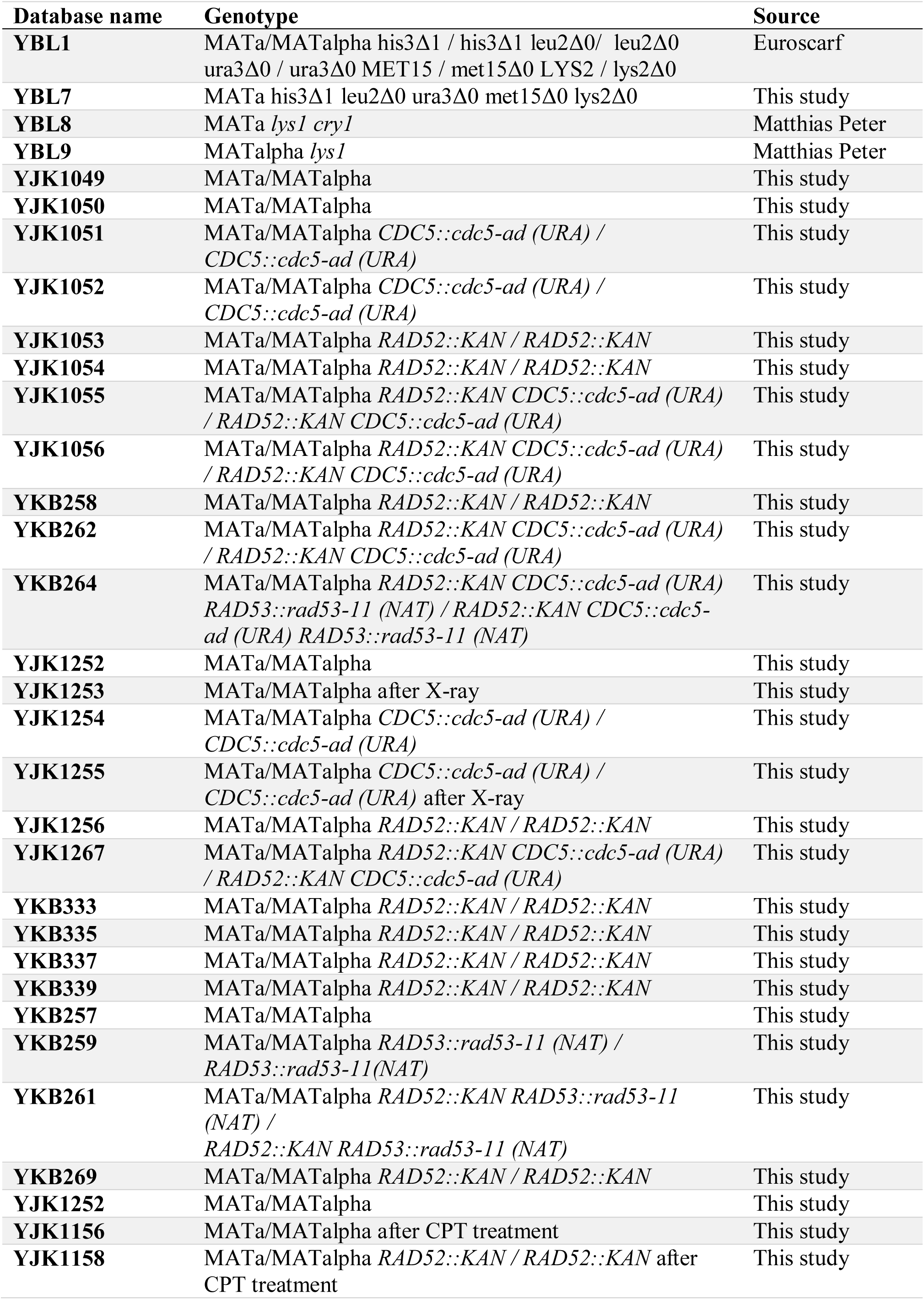
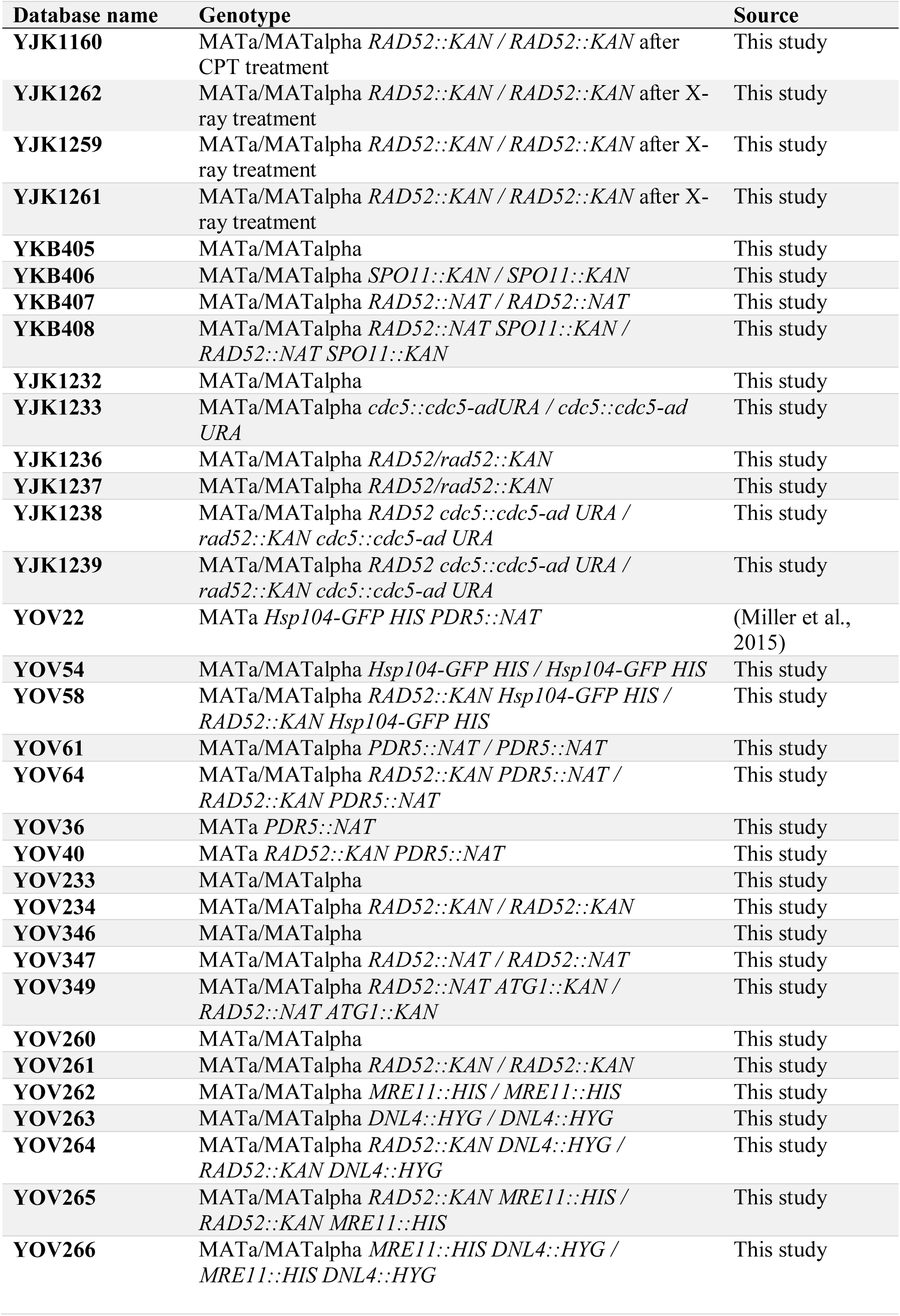
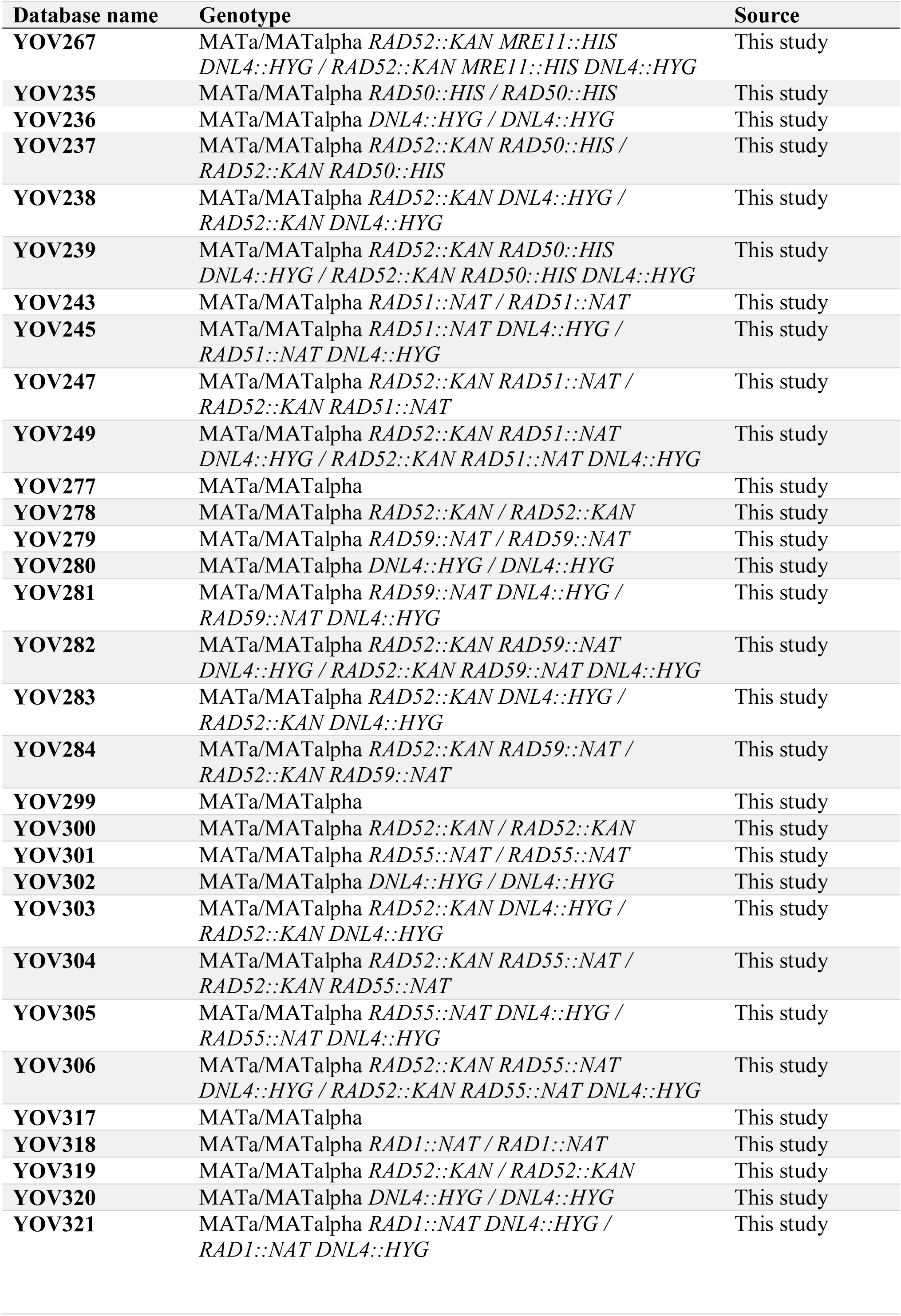
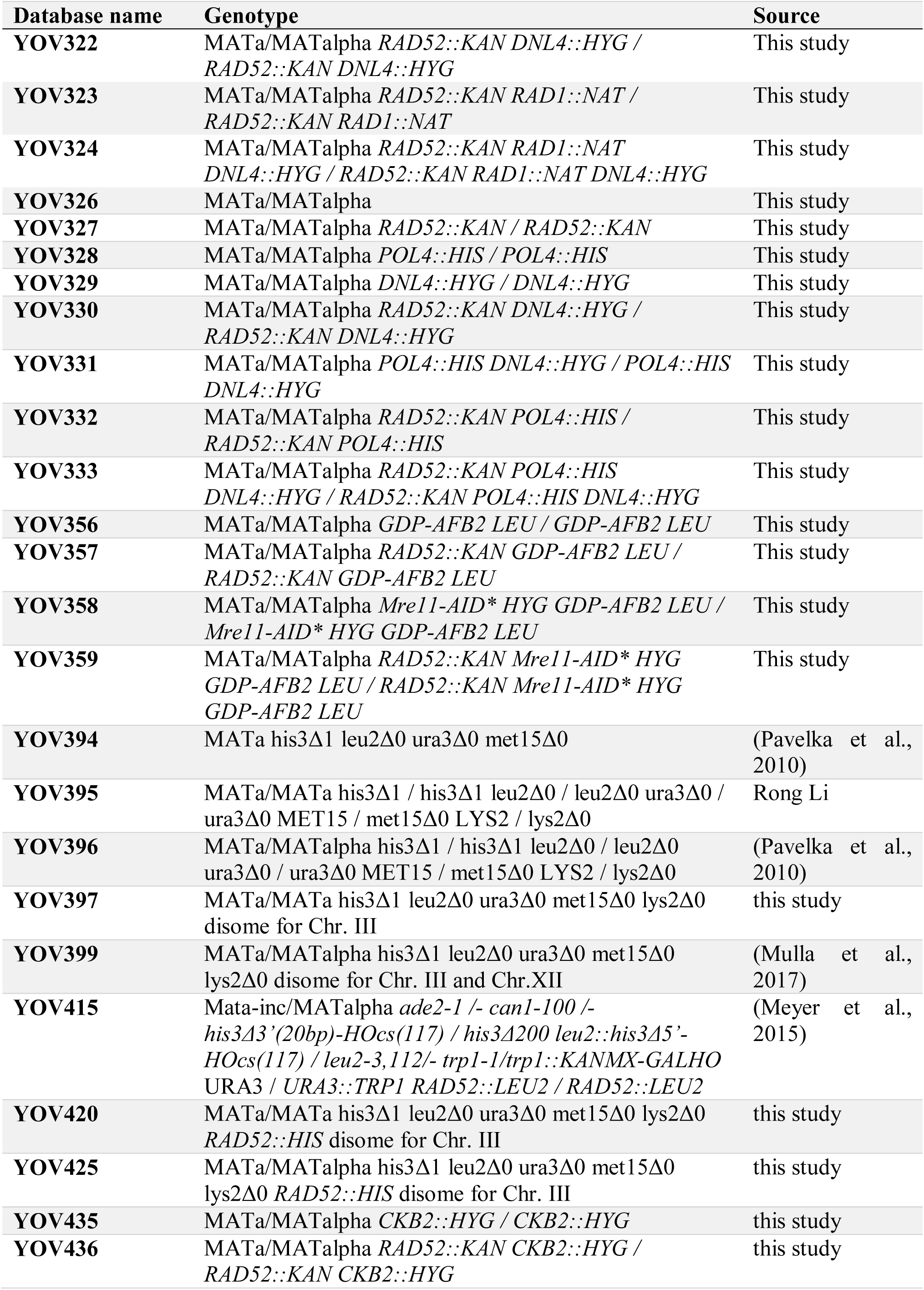
List of yeast strains used in this study

| Database name | Genotype | Source |
| --- | --- | --- |
| **YBL1** | MATa/MATalpha his3Δ1 / his3Δ1 leu2Δ0/ leu2Δ0<br>ura3Δ0 / ura3Δ0 MET15 / met15Δ0 LYS2 / lys2Δ0 | Euroscarf |
| **YBL7** | MATa his3Δ1 leu2Δ0 ura3Δ0 met15Δ0 lys2Δ0 | This study |
| **YBL8** | MATa *lys1 cry1* | Matthias Peter |
| **YBL9** | MATalpha *lys1* | Matthias Peter |
| **YJK1049** | MATa/MATalpha | This study |
| **YJK1050** | MATa/MATalpha | This study |
| **YJK1051** | MATa/MATalpha *CDC5::cdc5-ad (URA)* /<br>*CDC5::cdc5-ad (URA)* | This study |
| **YJK1052** | MATa/MATalpha *CDC5::cdc5-ad (URA)* /<br>*CDC5::cdc5-ad (URA)* | This study |
| **YJK1053** | MATa/MATalpha *RAD52::KAN / RAD52::KAN* | This study |
| **YJK1054** | MATa/MATalpha *RAD52::KAN / RAD52::KAN* | This study |
| **YJK1055** | MATa/MATalpha *RAD52::KAN CDC5::cdc5-ad (URA)*<br>/*RAD52::KAN CDC5::cdc5-ad (URA)* | This study |
| **YJK1056** | MATa/MATalpha *RAD52::KAN CDC5::cdc5-ad (URA)*<br>/*RAD52::KAN CDC5::cdc5-ad (URA)* | This study |
| **YKB258** | MATa/MATalpha *RAD52::KAN / RAD52::KAN* | This study |
| **YKB262** | MATa/MATalpha *RAD52::KAN CDC5::cdc5-ad (URA)*<br>/*RAD52::KAN CDC5::cdc5-ad (URA)* | This study |
| **YKB264** | MATa/MATalpha *RAD52::KAN CDC5::cdc5-ad (URA)*<br>*RAD53::rad53-11 (NAT) / RAD52::KAN CDC5::cdc5-*<br>*ad (URA) RAD53::rad53-11 (NAT)* | This study |
| **YJK1252** | MATa/MATalpha | This study |
| **YJK1253** | MATa/MATalpha after X-ray | This study |
| **YJK1254** | MATa/MATalpha *CDC5::cdc5-ad (URA)* /<br>*CDC5::cdc5-ad (URA)* | This study |
| **YJK1255** | MATa/MATalpha *CDC5::cdc5-ad (URA)* /<br>*CDC5::cdc5-ad (URA)* after X-ray | This study |
| **YJK1256** | MATa/MATalpha *RAD52::KAN / RAD52::KAN* | This study |
| **YJK1267** | MATa/MATalpha *RAD52::KAN CDC5::cdc5-ad (URA)*<br>/*RAD52::KAN CDC5::cdc5-ad (URA)* | This study |
| **YKB333** | MATa/MATalpha *RAD52::KAN / RAD52::KAN* | This study |
| **YKB335** | MATa/MATalpha *RAD52::KAN / RAD52::KAN* | This study |
| **YKB337** | MATa/MATalpha *RAD52::KAN / RAD52::KAN* | This study |
| **YKB339** | MATa/MATalpha *RAD52::KAN / RAD52::KAN* | This study |
| **YKB257** | MATa/MATalpha | This study |
| **YKB259** | MATa/MATalpha *RAD53::rad53-11 (NAT) /*<br>*RAD53::rad53-11(NAT)* | This study |
| **YKB261** | MATa/MATalpha *RAD52::KAN RAD53::rad53-11*<br>*(NAT) /*<br>*RAD52::KAN RAD53::rad53-11 (NAT)* | This study |
| **YKB269** | MATa/MATalpha *RAD52::KAN / RAD52::KAN* | This study |
| **YJK1252** | MATa/MATalpha | This study |
| **YJK1156** | MATa/MATalpha after CPT treatment | This study |
| **YJK1158** | MATa/MATalpha *RAD52::KAN / RAD52::KAN* after<br>CPT treatment | This study |
| YJK1160 | MATa/MATalpha *RAD52::KAN / RAD52::KAN* after CPT treatment | This study |
| YJK1262 | MATa/MATalpha *RAD52::KAN / RAD52::KAN* after X-ray treatment | This study |
| YJK1259 | MATa/MATalpha *RAD52::KAN / RAD52::KAN* after X-ray treatment | This study |
| YJK1261 | MATa/MATalpha *RAD52::KAN / RAD52::KAN* after X-ray treatment | This study |
| YKB405 | MATa/MATalpha | This study |
| YKB406 | MATa/MATalpha *SPO11::KAN / SPO11::KAN* | This study |
| YKB407 | MATa/MATalpha *RAD52::NAT / RAD52::NAT* | This study |
| YKB408 | MATa/MATalpha *RAD52::NAT SPO11::KAN / RAD52::NAT SPO11::KAN* | This study |
| YJK1232 | MATa/MATalpha | This study |
| YJK1233 | MATa/MATalpha *cdc5::cdc5-adURA / cdc5::cdc5-adURA* | This study |
| YJK1236 | MATa/MATalpha *RAD52/rad52::KAN* | This study |
| YJK1237 | MATa/MATalpha *RAD52/rad52::KAN* | This study |
| YJK1238 | MATa/MATalpha *RAD52 cdc5::cdc5-ad URA / rad52::KAN cdc5::cdc5-ad URA* | This study |
| YJK1239 | MATa/MATalpha *RAD52 cdc5::cdc5-ad URA / rad52::KAN cdc5::cdc5-ad URA* | This study |
| YOV22 | MATa *Hsp104-GFP HIS PDR5::NAT* | (Miller et al., 2015) |
| YOV54 | MATa/MATalpha *Hsp104-GFP HIS / Hsp104-GFP HIS* | This study |
| YOV58 | MATa/MATalpha *RAD52::KAN Hsp104-GFP HIS / RAD52::KAN Hsp104-GFP HIS* | This study |
| YOV61 | MATa/MATalpha *PDR5::NAT / PDR5::NAT* | This study |
| YOV64 | MATa/MATalpha *RAD52::KAN PDR5::NAT / RAD52::KAN PDR5::NAT* | This study |
| YOV36 | MATa *PDR5::NAT* | This study |
| YOV40 | MATa *RAD52::KAN PDR5::NAT* | This study |
| YOV233 | MATa/MATalpha | This study |
| YOV234 | MATa/MATalpha *RAD52::KAN / RAD52::KAN* | This study |
| YOV346 | MATa/MATalpha | This study |
| YOV347 | MATa/MATalpha *RAD52::NAT / RAD52::NAT* | This study |
| YOV349 | MATa/MATalpha *RAD52::NAT ATG1::KAN / RAD52::NAT ATG1::KAN* | This study |
| YOV260 | MATa/MATalpha | This study |
| YOV261 | MATa/MATalpha *RAD52::KAN / RAD52::KAN* | This study |
| YOV262 | MATa/MATalpha *MRE11::HIS / MRE11::HIS* | This study |
| YOV263 | MATa/MATalpha *DNL4::HYG / DNL4::HYG* | This study |
| YOV264 | MATa/MATalpha *RAD52::KAN DNL4::HYG / RAD52::KAN DNL4::HYG* | This study |
| YOV265 | MATa/MATalpha *RAD52::KAN MRE11::HIS / RAD52::KAN MRE11::HIS* | This study |
| YOV266 | MATa/MATalpha *MRE11::HIS DNL4::HYG / MRE11::HIS DNL4::HYG* | This study |

| **Database name** | **Genotype** | **Source** |
| --- | --- | --- |
| **YOV267** | MATa/MATalpha *RAD52::KAN MRE11::HIS*<br>*DNL4::HYG / RAD52::KAN MRE11::HIS DNL4::HYG* | This study |
| **YOV235** | MATa/MATalpha *RAD50::HIS / RAD50::HIS* | This study |
| **YOV236** | MATa/MATalpha *DNL4::HYG / DNL4::HYG* | This study |
| **YOV237** | MATa/MATalpha *RAD52::KAN RAD50::HIS /*<br>*RAD52::KAN RAD50::HIS* | This study |
| **YOV238** | MATa/MATalpha *RAD52::KAN DNL4::HYG /*<br>*RAD52::KAN DNL4::HYG* | This study |
| **YOV239** | MATa/MATalpha *RAD52::KAN RAD50::HIS*<br>*DNL4::HYG / RAD52::KAN RAD50::HIS DNL4::HYG* | This study |
| **YOV243** | MATa/MATalpha *RAD51::NAT / RAD51::NAT* | This study |
| **YOV245** | MATa/MATalpha *RAD51::NAT DNL4::HYG /*<br>*RAD51::NAT DNL4::HYG* | This study |
| **YOV247** | MATa/MATalpha *RAD52::KAN RAD51::NAT /*<br>*RAD52::KAN RAD51::NAT* | This study |
| **YOV249** | MATa/MATalpha *RAD52::KAN RAD51::NAT*<br>*DNL4::HYG / RAD52::KAN RAD51::NAT DNL4::HYG* | This study |
| **YOV277** | MATa/MATalpha | This study |
| **YOV278** | MATa/MATalpha *RAD52::KAN / RAD52::KAN* | This study |
| **YOV279** | MATa/MATalpha *RAD59::NAT / RAD59::NAT* | This study |
| **YOV280** | MATa/MATalpha *DNL4::HYG / DNL4::HYG* | This study |
| **YOV281** | MATa/MATalpha *RAD59::NAT DNL4::HYG /*<br>*RAD59::NAT DNL4::HYG* | This study |
| **YOV282** | MATa/MATalpha *RAD52::KAN RAD59::NAT*<br>*DNL4::HYG / RAD52::KAN RAD59::NAT DNL4::HYG* | This study |
| **YOV283** | MATa/MATalpha *RAD52::KAN DNL4::HYG /*<br>*RAD52::KAN DNL4::HYG* | This study |
| **YOV284** | MATa/MATalpha *RAD52::KAN RAD59::NAT /*<br>*RAD52::KAN RAD59::NAT* | This study |
| **YOV299** | MATa/MATalpha | This study |
| **YOV300** | MATa/MATalpha *RAD52::KAN / RAD52::KAN* | This study |
| **YOV301** | MATa/MATalpha *RAD55::NAT / RAD55::NAT* | This study |
| **YOV302** | MATa/MATalpha *DNL4::HYG / DNL4::HYG* | This study |
| **YOV303** | MATa/MATalpha *RAD52::KAN DNL4::HYG /*<br>*RAD52::KAN DNL4::HYG* | This study |
| **YOV304** | MATa/MATalpha *RAD52::KAN RAD55::NAT /*<br>*RAD52::KAN RAD55::NAT* | This study |
| **YOV305** | MATa/MATalpha *RAD55::NAT DNL4::HYG /*<br>*RAD55::NAT DNL4::HYG* | This study |
| **YOV306** | MATa/MATalpha *RAD52::KAN RAD55::NAT*<br>*DNL4::HYG / RAD52::KAN RAD55::NAT DNL4::HYG* | This study |
| **YOV317** | MATa/MATalpha | This study |
| **YOV318** | MATa/MATalpha *RAD1::NAT / RAD1::NAT* | This study |
| **YOV319** | MATa/MATalpha *RAD52::KAN / RAD52::KAN* | This study |
| **YOV320** | MATa/MATalpha *DNL4::HYG / DNL4::HYG* | This study |
| **YOV321** | MATa/MATalpha *RAD1::NAT DNL4::HYG /*<br>*RAD1::NAT DNL4::HYG* | This study |
| YOV322 | MATa/MATalpha *RAD52::KAN DNL4::HYG* / *RAD52::KAN DNL4::HYG* | This study |
| YOV323 | MATa/MATalpha *RAD52::KAN RAD1::NAT* / *RAD52::KAN RAD1::NAT* | This study |
| YOV324 | MATa/MATalpha *RAD52::KAN RAD1::NAT DNL4::HYG* / *RAD52::KAN RAD1::NAT DNL4::HYG* | This study |
| YOV326 | MATa/MATalpha | This study |
| YOV327 | MATa/MATalpha *RAD52::KAN* / *RAD52::KAN* | This study |
| YOV328 | MATa/MATalpha *POL4::HIS* / *POL4::HIS* | This study |
| YOV329 | MATa/MATalpha *DNL4::HYG* / *DNL4::HYG* | This study |
| YOV330 | MATa/MATalpha *RAD52::KAN DNL4::HYG* / *RAD52::KAN DNL4::HYG* | This study |
| YOV331 | MATa/MATalpha *POL4::HIS DNL4::HYG* / *POL4::HIS DNL4::HYG* | This study |
| YOV332 | MATa/MATalpha *RAD52::KAN POL4::HIS* / *RAD52::KAN POL4::HIS* | This study |
| YOV333 | MATa/MATalpha *RAD52::KAN POL4::HIS DNL4::HYG* / *RAD52::KAN POL4::HIS DNL4::HYG* | This study |
| YOV356 | MATa/MATalpha *GDP-AFB2 LEU* / *GDP-AFB2 LEU* | This study |
| YOV357 | MATa/MATalpha *RAD52::KAN GDP-AFB2 LEU* / *RAD52::KAN GDP-AFB2 LEU* | This study |
| YOV358 | MATa/MATalpha *Mre11-AID\* HYG GDP-AFB2 LEU* / *Mre11-AID\* HYG GDP-AFB2 LEU* | This study |
| YOV359 | MATa/MATalpha *RAD52::KAN Mre11-AID\* HYG GDP-AFB2 LEU* / *RAD52::KAN Mre11-AID\* HYG GDP-AFB2 LEU* | This study |
| YOV394 | MATa his3Δ1 leu2Δ0 ura3Δ0 met15Δ0 | (Pavelka et al., 2010) |
| YOV395 | MATa/MATa his3Δ1 / his3Δ1 leu2Δ0 / leu2Δ0 ura3Δ0 / ura3Δ0 MET15 / met15Δ0 LYS2 / lys2Δ0 | Rong Li |
| YOV396 | MATa/MATalpha his3Δ1 / his3Δ1 leu2Δ0 / leu2Δ0 ura3Δ0 / ura3Δ0 MET15 / met15Δ0 LYS2 / lys2Δ0 | (Pavelka et al., 2010) |
| YOV397 | MATa/MATa his3Δ1 leu2Δ0 ura3Δ0 met15Δ0 lys2Δ0 disome for Chr. III | this study |
| YOV399 | MATa/MATalpha his3Δ1 leu2Δ0 ura3Δ0 met15Δ0 lys2Δ0 disome for Chr. III and Chr. XII | (Mulla et al., 2017) |
| YOV415 | Mata-inc/MATalpha *ade2-1* /- *can1-100* /- *his3Δ3'(20bp)-HOcs(117)* / *his3Δ200 leu2::his3Δ5'-HOcs(117)* / *leu2-3,112/- trp1-1/trp1::KANMX-GALHO URA3* / *URA3::TRP1 RAD52::LEU2* / *RAD52::LEU2* | (Meyer et al., 2015) |
| YOV420 | MATa/MATa his3Δ1 leu2Δ0 ura3Δ0 met15Δ0 lys2Δ0 *RAD52::HIS* disome for Chr. III | this study |
| YOV425 | MATa/MATalpha his3Δ1 leu2Δ0 ura3Δ0 met15Δ0 lys2Δ0 *RAD52::HIS* disome for Chr. III | this study |
| YOV435 | MATa/MATalpha *CKB2::HYG* / *CKB2::HYG* | this study |
| YOV436 | MATa/MATalpha *RAD52::KAN CKB2::HYG* / *RAD52::KAN CKB2::HYG* | this study |

**Supplementary Table 2.**
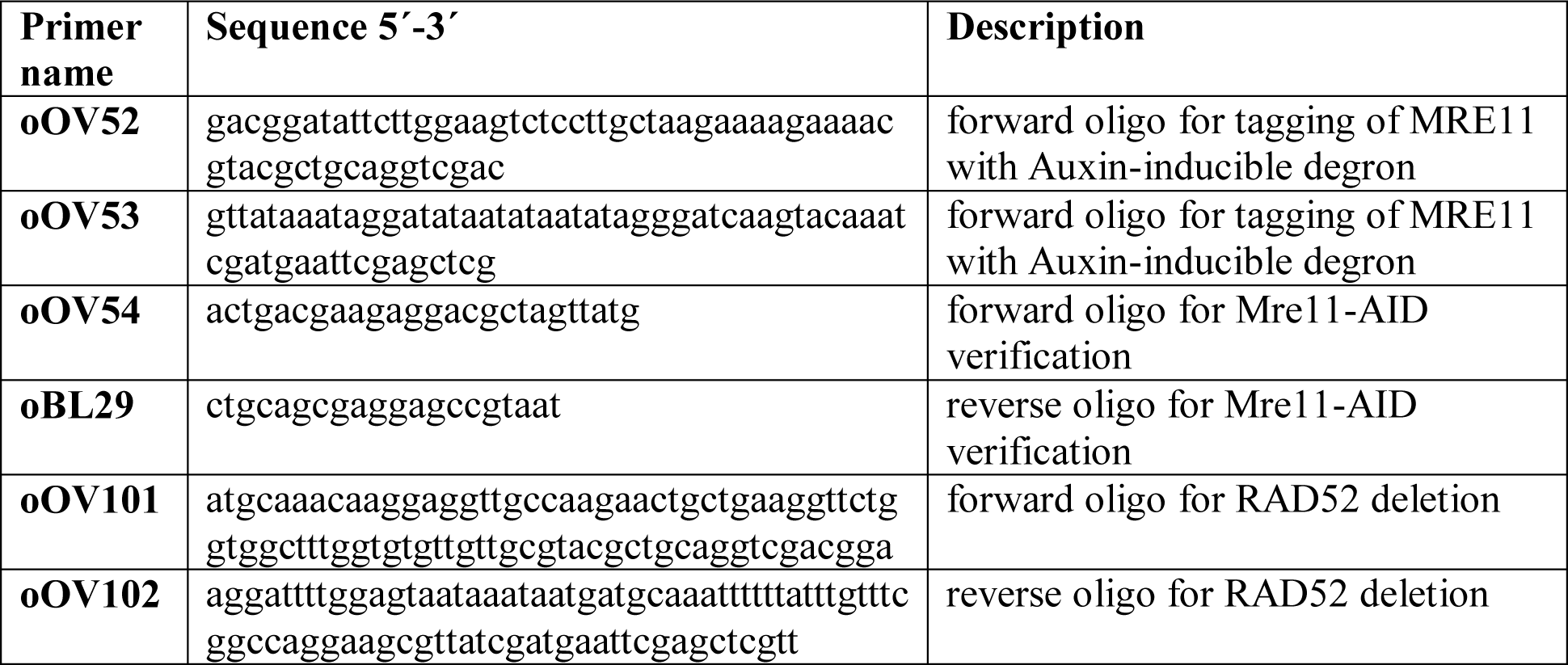
List of primers used in this study

**Supplementary Table 3.** List of plasmids used in this study

| **Plasmid name** | **Description** | **Source** |
| --- | --- | --- |
| pBL186 | pRS423-GAL-HIS3, empty vector | (Mumberg et al., 1994, 1995) |
| pBL211 | pRS425-GAL-LEU2, empty vector | (Mumberg et al., 1994, 1995) |
| pBL299 | pRS41-hygB, empty vector | (Taxis and Knop, 2006) |
| pBL303 | pRS42-KAN, empty vector | (Taxis and Knop, 2006) |
| pBL304 | pRS42-NAT, empty vector | (Taxis and Knop, 2006) |
| pJK34 | pRS315-GFP-ATG8 | (Klermund et al., 2014) |
| pBL454 | pSM409-pHyg-AID*-6FLAG | (Morawska and Ulrich, 2013) |
| pBL252 | pFA6A-HIS3MX6 | (Longtine et al., 1998) |

**Supplementary Table 4.** List of antibodies used in this study

| **Antibodies** | **Dilution** | **Source** |
| --- | --- | --- |
| **Primary antibodies** |  |  |
| mouse anti-Rad53 EL7.E1 | 1:16 | Gift from M. Foiani |
| mouse anti-Rad53 EL7.E1 | 1:1000 | Abcam, ab166859 |
| mouse anti-Pgk1 | 1:200 000 | Invitrogen, #459250 |
| rabbit anti-Rnr3 | 1:300 | Agrisera, #AS09 574 |
| mouse anti-actin | 1:2000 | Millipore, MAB1501R |
| rabbit anti-tubulin | 1:3000 | Abcam, ab184970 |
| rabbit anti-Sch9 | 1:20000 | Gift from R. Loewith |
| mouse anti-phospho Sch9 | 1:10000 | Gift from R. Loewith |
| mouse anti-GFP | 1:1000 | Roche Diagnostics via Sigma-Aldrich,<br># 11 814 460 001 |
| **Secondary antibodies** |  |  |
| goat anti-mouse (HRP conjugate) | 1:3000 | Bio-Rad, #170-5047 |
| goat anti-rabbit (HRP conjugate) | 1:3000 | Bio-Rad, #170-5046 |
| goat anti-rabbit and anti-mouse<br>IRDye 680 | 1:10 000 | Licor, #926-68071 and #926-68070 |
| goat anti-rabbit and anti-mouse<br>IRDye 800 | 1:10 000 | Licor, #926-32211 and #926-32210 |

